# Fructose-1-kinase has pleiotropic roles in *Escherichia coli*

**DOI:** 10.1101/2023.12.14.571569

**Authors:** Chamitha Weeramange, Cindy Menjivar, Pierce T. O’Neil, Samir El Qaidi, Kelly S. Harrison, Sarah Meinhardt, Cole L. Bird, Shwetha Sreenivasan, Philip R. Hardwidge, Aron W. Fenton, P. Scott Hefty, Jeffrey L. Bose, Liskin Swint-Kruse

## Abstract

In *Escherichia coli*, the master transcription regulator Catabolite Repressor Activator (Cra) regulates >100 genes in central metabolism. Cra binding to DNA is allosterically regulated by binding to fructose-1-phosphate (F-1-P), but the only documented source of F-1-P is from the concurrent import and phosphorylation of exogenous fructose. Thus, many have proposed that fructose-1,6-bisphosphate (F-1,6-BP) is also a physiological regulatory ligand. However, the role of F-1,6-BP has been widely debated. Here, we report that the *E. coli* enzyme fructose-1-kinase (FruK) can carry out its “reverse” reaction under physiological substrate concentrations to generate F-1-P from F-1,6-BP. We further show that FruK directly binds Cra with nanomolar affinity and forms higher order, heterocomplexes. Growth assays with a Δ*fruK* strain and *fruK* complementation show that FruK has a broader role in metabolism than fructose catabolism. The Δ*fruK* strain also alters biofilm formation. Since *fruK* itself is repressed by Cra, these newly-reported events add layers to the dynamic regulation of *E. coli* central metabolism that occur in response to changing nutrients. These findings might have wide-spread relevance to other γ-proteobacteria, which conserve both Cra and FruK.

## Introduction

The ability of a microorganism to respond to external stimuli, such as changes in available carbon sources, requires the ability to switch between alternative metabolic programs. Many switches are accomplished by the activity of transcription factors that control the expression of different groups of metabolic genes. In γ-proteobacteria, one of the most important transcription factors is the Catabolite Repressor Activator protein (Cra; UniProt P0ACP1). In *Escherichia coli*, Cra differentially regulates genes important to glycolysis, TCA cycle, gluconeogenesis, and the Entner-Doudorhoff pathway^1–6^ so that glucose metabolism is repressed when gluconeogenesis is activated. A key step of this regulatory process occurs when Cra binds fructose metabolite(s) that, in turn, allosterically diminish Cra’s binding to DNA^7^.

An unambiguous finding is that fructose-1-phosphate (F-1-P) can act as a strong Cra allosteric inhibitor at micromolar concentrations^7^. However, the biological source(s) of F-1-P and whether it is the only fructose metabolite that allosterically regulates Cra has engendered some controversy. To date, the formation of F-1-P has only been documented as arising from the coincident import and phosphorylation of exogenous fructose by FruBA (Figure 1; ^8–12^). However, it is something of a paradox that F-1-P from exogenous fructose would serve as the only molecular switch for this central metabolic network. Indeed, results from a variety of biological studies (*e.g.,* ^13–17^), are consistent with the proposal that millimolar concentrations of fructose-1,6-bisphosphate (F-1,6-BP) also regulate Cra’s activity^7^. Since F-1,6-BP is a metabolic intermediate for many sugars, this would allow Cra to be regulated in a wide variety of nutrient conditions.

**Figure 1.**
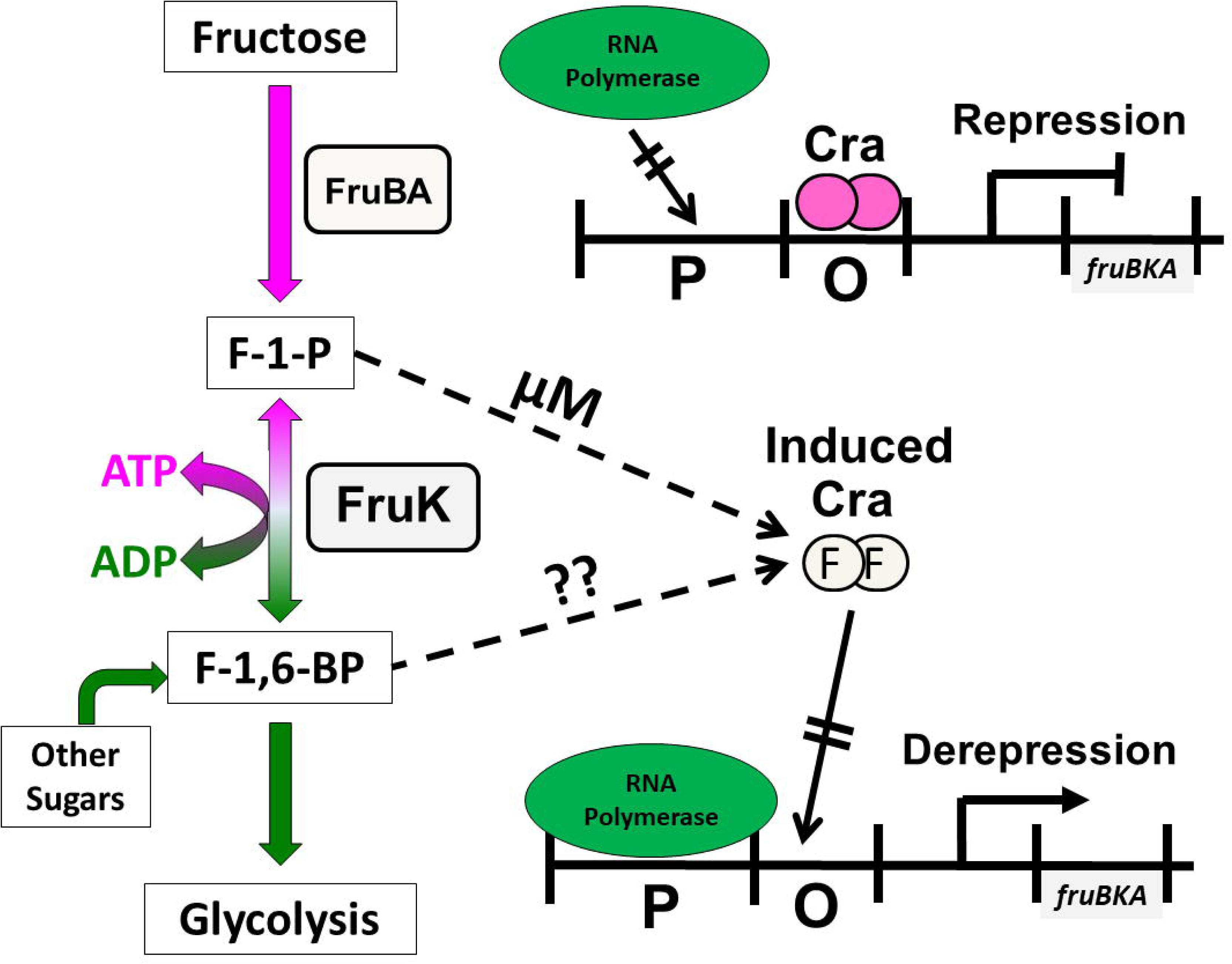
Known co-regulation of Cra and FruK. The left portion of the cartoon shows the relationship and source(s) of F-1-P and F-1,6-BP. F-1-P can be formed by the concurrent import and phosphorylation of fructose by FruBA. F-1,6-BP can be produced from many other sugars that funnel into glycolysis. The enzyme FruK was previously known to phosphorylate F-1-P to F-1,6-BP; in this work, we show that the reverse reaction can also occur. Transcriptional regulation of these fructose metabolic enzymes occurs when Cra binds to the operator of the *fruBKA* operon and represses downstream genes (upper left). When micromolar concentrations of F-1-P are present, F-1-P binds to Cra and to diminishes Cra’s binding affinity for operator DNA, thereby inducing transcription of the *fruBKA* genes (lower right). The direct effect of F-1,6-BP binding to Cra has been disputed^20^, leading to a conundrum about the biological relationship observed between F-1,6-BP levels and Cra regulatory activities^13–17^.

However, there is limited evidence for direct F-1,6-BP induction of the Cra regulon: In an early study, the effect of 5 mM F-1,6-BP on Cra’s DNA binding was weak; the authors acknowledged that the effect could be due to F-1-P contamination^7^. Study of the *Pseudomonas putida* Cra supported the contamination hypothesis^18,19^. Most convincingly, a 2018 study of *E. coli* Cra showed no direct F-1,6-BP binding or induction from the *fruB_distal_* DNA operator binding site^20^.

Efforts to experimentally assess and model the Cra regulon have not helped to clarify the role of F-1,6-BP or the source of F-1-P: In one study, Cra was proposed to act as a flux sensor that uses intracellular F-1,6-BP to integrate information from a variety of nutrients^16^. However, most of the *in vivo* F-1,6-BP concentrations measured under the study conditions were less than 1 mM, nearly an order of magnitude lower than that needed for this metabolite to weakly regulate Cra^7^ (if F-1,6-BP can even regulate Cra^20^). A second study to model *E. coli* metabolic flux used F-1-P as the only Cra modulator, but the model predictions could not be reconciled with experimental evidence^21^. A third study found that F-1,6-BP, F-1-P, and cyclic AMP were the three key metabolites that regulate central metabolism, with F-1-P and F-1,6-BP presumably acting through their interaction with Cra^22^. Indeed, among 47 central metabolites measured across 23 different growth conditions, F-1-P displayed the highest variance in intracellular concentration, and its concentration generally correlated with F-1,6-BP concentration (Supplemental Figure 1; adapted from ^22^). Since exogenous fructose was only provided for one of the 23 growth conditions, the detected F-1-P must be produced in *E. coli* from other sources.

One way to reconcile these observations would be if F-1,6-BP is converted to F-1-P. An obvious, although largely overlooked, means by which this could occur is via the “reverse” reaction of fructose-1-kinase (FruK; a member of ribokinase family^23^; UniProt P0AEW9)^11,24^. To date, the only documented role for FruK is its “forward” reaction, which occurs after import/phosphorylation of exogenous fructose and uses ATP to convert F-1-P to F-1,6-BP and ADP (Supplemental Figure 2). However, all enzymes have the potential to catalyze their chemical reactions in both directions. Thus, we hypothesized that, if the reverse FruK reaction can occur under physiological concentrations (Supplemental Table 1), then FruK can affect Cra’s function by using F-1,6-BP derived from other carbon sources to generate F-1-P. Another intriguing possibility is that, since FruK was observed to directly interact with a chimeric protein containing the Cra regulatory domain (“LLhF”; Supplemental Figure 3)^25,26^, FruK might also directly interact with wild-type Cra and impact its activity.

In the study described herein, we assessed whether FruK carries out its reverse reaction, the potential for direct interactions between FruK and Cra, and contributions of FruK to *E. coli* metabolism. Our results suggest that FruK plays pleiotropic roles in *E. coli* that contribute to its bacterial physiology.

## Results

### FruK can generate F-1-P from physiological concentrations of F-1,6-BP and ADP

The physiological concentrations of F-1,6-BP, ADP, and ATP have been measured under a variety of growth conditions in at least two prior studies (Supplemental Table 1)^22,27^. These data are replotted in Supplemental Figure 1 to show the relationship between F-1-P, and F-1-6-BP. When *E. coli* are grown in the presence of glucose, F-1,6-BP can be as high as 15 mM^27^. To determine whether FruK can use F-1,6-BP to generate Cra’s strong allosteric regulator F-1-P, we assessed whether this enzyme can carry out its “reverse” reaction (Figure 1) at 3 mM ADP and 0.1-15 mM FBP.

Since the reverse reaction generates ATP, we monitored the reaction by coupling ATP formation to a luciferase assay. ATP generation was detectable in a concentration-dependent manner across a range of physiological F-1,6-BP substrate concentrations (Figure 2) demonstrating that FruK can indeed generate F-1-P from F-1,6-BP. Since we hypothesized that Cra and FruK can also directly form a protein-protein complex (described below), we tested if the presence of Cra would impact FruK activity. The addition of Cra, with or without DNA, had no significant impact on FruK activity.

**Figure 2.**
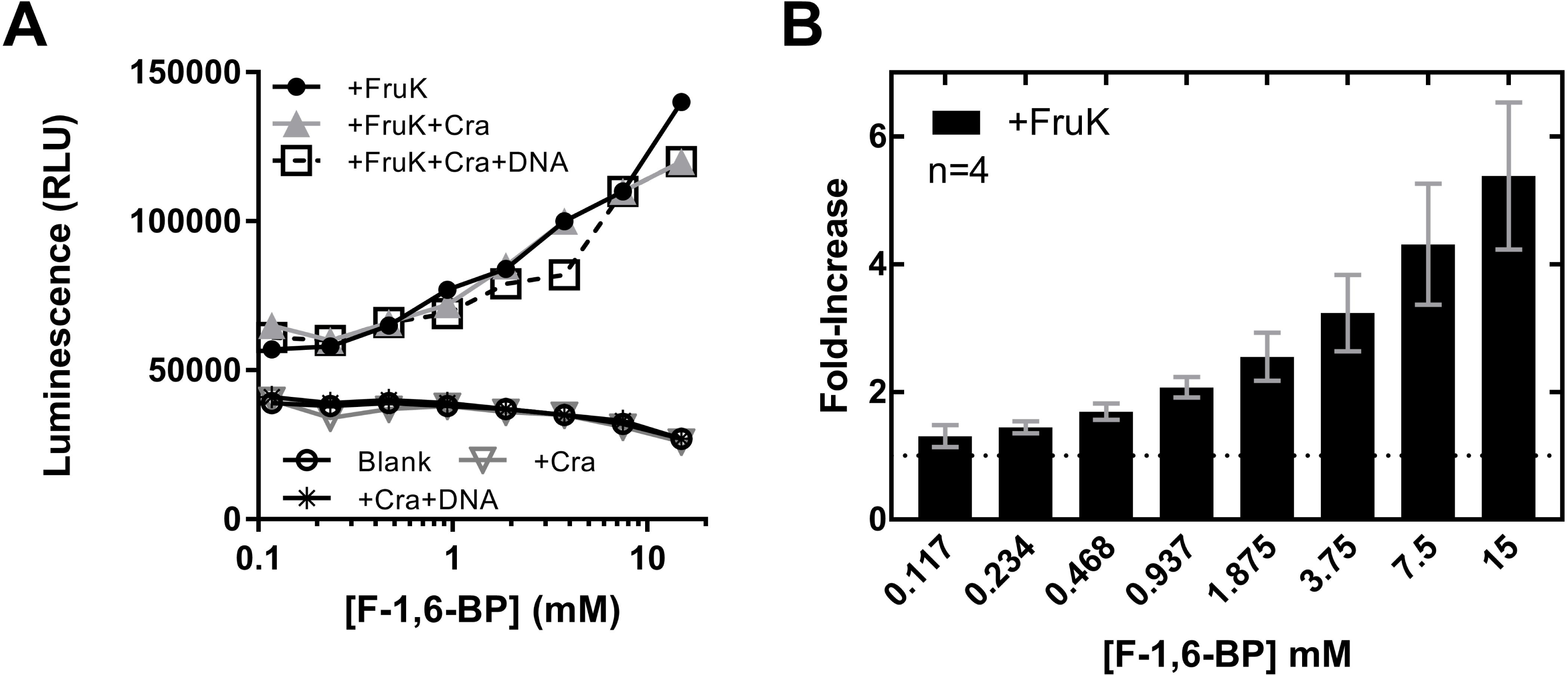
FruK generates ATP from ADP and F-1,6-BP. The “reverse” FruK reaction F-1,6-BP + ADP E- -+ F-1-P + ATP was coupled to a luciferase assay to detect generation of ATP. The assays shown in these panels maintained constant concentrations of counter ions across the F-1,6-BP range. (A) Representative raw data for the luciferase reaction in the presence and absence of FruK, Cra, and DNA; all experiments in this panel were repeated at least two independent times; the same result was obtained across a wide variety of buffer conditions and protein concentrations in multiple additional independent experiments. In the experiments shown here, both FruK and Cra were 275 nM. DNA (*fruB_distal_*) was 30 nM; although this DNA concentration did not saturate Cra, raising the DNA concentration further would increase the ionic strength. Supporting assays (Supplemental Figure 9) showed that increasing ion strength inhibited the luciferase reaction. (B) The average fold-increase was calculated from 4 independent replicates of the +FruK luciferase assay at different F-1,6-BP concentrations, after normalizing to the buffer-only control; error bars represent the standard deviations of the averages.

### FruK binds wild-type Cra in E. coli crude cell extracts

Our attention was first called to FruK in prior studies that used DNA pull-down assays to assess the functions of chimeric LacI/GalR homologs^26^. In that study, FruK interacted with the LacI-Cra chimera “LLhF”, which was composed of the DNA binding domain of the *E. coli* lactose repressor protein (LacI) and the Cra regulatory domain (domain organization is illustrated in Supplemental Figure 3) but not with full-length LacI or other chimeras^26^. This finding suggested that FruK could interact with the regulatory domain of wild-type Cra.

To test for the potential interaction between wild-type Cra and FruK, we again used a DNA pull-down assay. First, as a positive control, we repeated the experiment in which LLhF was over-expressed from a constitutive promoter and confirmed that it did indeed pull-down FruK when using immobilized *lacO*^1^ operator^28^ as a bait (Figure 3A, lane 1). Next, we adapted the pull-down assay to use the consensus Cra DNA binding site (*O^cra^*)^2^ as bait for constitutively over-expressed Cra. In this case, a band for Cra was readily identified, but no FruK was detected (Figure 3A, lane 3). However, when fructose was added to the growth media (which would induce transcription of the *fruBKA* operon and thereby increase levels FruK protein^7^), a band for FruK was observed along with Cra (Figure 3A, lane 6). Possible explanations for why LLhF does not require fructose in induce high levels of the FruK protein are discussed in Supplemental Results. As negative controls, pull-down samples from strains that did not constitutively over-express Cra showed no Cra or FruK in either the absence or presence of fructose (Figure 3A, lanes 5 and 7). Together, these data support the hypothesis that FruK binds directly to Cra.

**Figure 3.**
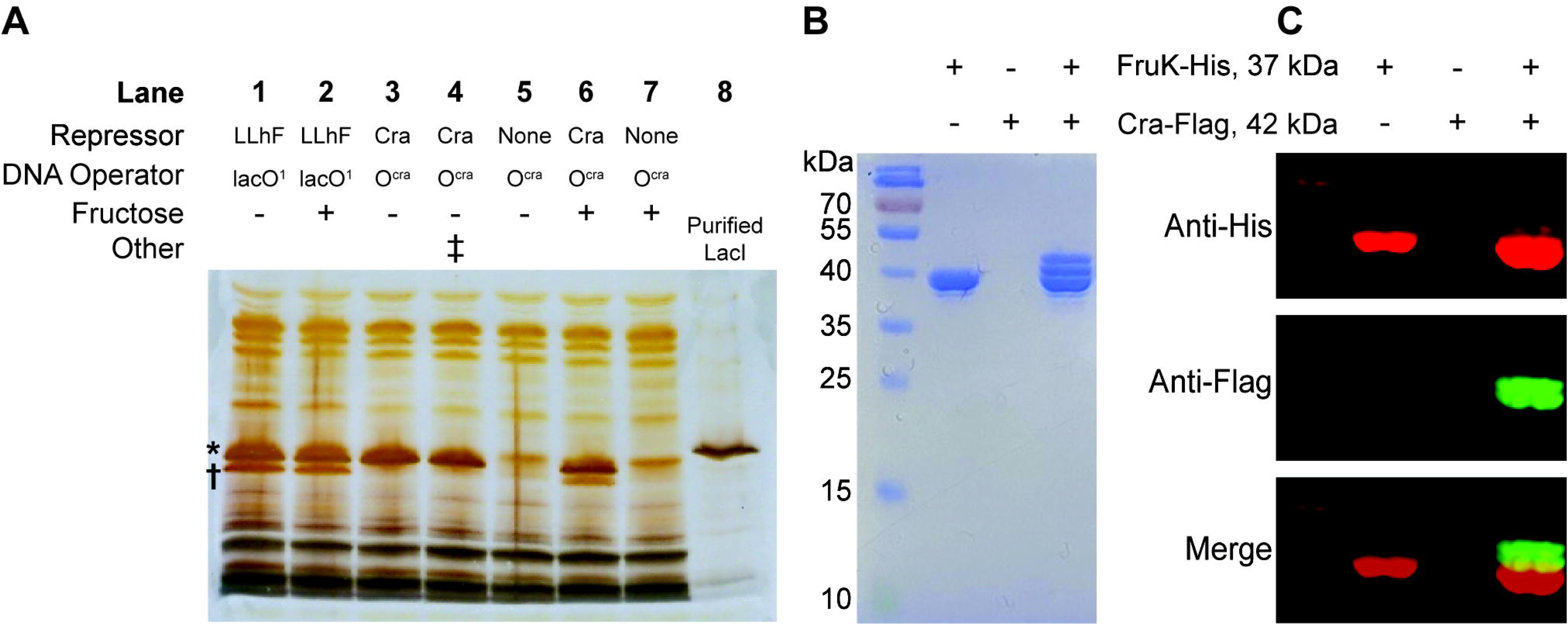
Cra forms complexes with FruK in *E. coli* crude cell extracts. (A) DNA pull-down assays using exogenous Cra and LLhF chimera show that both proteins interact with endogenous FruK. LLhF (lanes 1-2) and wild-type Cra (lanes 3, 4 and 6) were constitutively over-expressed in *E. coli* 3.300 cells; control cells (lanes 5 and 7) lacked exogenous repressor. All FruK was expressed from the genomic *fruBKA* operon. Cells were grown in MOPS media with glycerol as the carbon source, in either the absence (lanes 1 and 3-5) or presence of 20 mM fructose (lanes 2 and 6-7). Lysed cells were incubated with the indicated immobilized DNA (lacO^1^ or O^cra^) and adherent proteins were visualized with SDS-PAGE and silver stain. The asterisk (*) indicates the position of the repressor (top) and the dagger (†) indicates the position of FruK (bottom); the identities of LLhF and FruK were previously ascertained with mass spectrometry.^25^ In lane 4, the double dagger (‡) indicates that lysates from lanes 3 and 5 were mixed prior to the pull-down assay. In lane 8, purified *E. coli* LacI serves as a molecular weight marker: LacI (360 a.a.; 38,590 Da monomer; UniProt P03023) is slightly larger than Cra (334 a.a.; 37,999 Da monomer; UniProt P0ACP1), which in turn is slightly larger than FruK (312 a.a.; 33,756 Da monomer; UniProt P0AEW9). This result was shown in at least two independent experiments. (B-C) Protein pull-down assays from crude cell extracts with over-expressed and tagged Cra and FruK in the absence of exogenous operator DNA. Cell lysates prepared from BL21 (DE3) *E. coli* expressing FruK-His were mixed with those expressing Cra-Flag, then subjected to a pull-down assay using nickel-agarose beads and SDS-PAGE. Results shown represent three independent experiments. Protein detection was by (B) Coomassie staining and (C) Western blot; the row headers of panels B and C are identical. Eluted proteins were immunoblotted using anti-His (red, top panel) and anti-Flag (green, middle panel) monoclonal antibodies. The lower panel is an overlay. Note that FruK-His pulls down Cra of two different sizes. The smaller band likely corresponds to Cra that has lost its N-terminal DNA binding domain (~50 amino acids). This domain is easily lost in Cra and other LacI/GalR proteins^94^, most likely to proteolysis^95–97^. The uncropped gels used to construct this figure are in Supplemental Figure 10.

In a complementary protein-protein pull-down experiment (*without* the addition of *O^cra^*), His-tagged FruK from enterohemorrhagic *E. coli* (EHEC) was over-expressed in one bacterial culture and Flag-tagged Cra was over-expressed in another. When the two cell lysates were mixed, only two proteins were detected in the nickel-resin elution (Figure 3B) and the identity of each band was confirmed by Western blot (Figure 3C). These results suggest that the Cra-FruK reaction is specific and can occur in the absence of DNA.

### FruK binds directly to Cra in vitro

Next, we directly examined the putative interaction between Cra and FruK using purified proteins and an adaptation of biolayer interferometry (“BLI”)^[25]^ to quantify the interaction. We previously used this technique to quantify binding between purified FruK and the DNA-bound LLhF chimera; the binding affinity was ~10^−8^ M ^[25]^. Also in this previously-published study, purified FruK did *not* bind to DNA-bound, purified LacI^25^, again supporting the hypothesis that FruK’s interaction with LLhF was mediated by its Cra regulatory domain rather than its LacI DNA binding domain.

Here, we performed BLI equilibrium assays with purified full-length Cra and FruK; data from an example experiment at one FruK concentration are shown in Supplemental Figure 4. In these experiments, biotinylated *O^cra^* DNA was immobilized to streptavidin fiber optic tips and then bound to purified Cra; this complex was then incubated with varied concentrations of FruK. Once equilibration was reached, the height of the signal plateau was plotted versus FruK concentration to generate titration curves (Figure 4)^25^. To calculate K_d_ from these curves, one must simultaneously determine the effective concentration of immobilized, DNA-bound Cra.^25^ This iterative process involved varying the amount of DNA-bound Cra that is immobilized on the tip, followed by global fitting of multiple independent experiments using the expanded equation for ligand binding (see Methods; Figure 4). This analysis showed that the assay conditions were in the “intermediate” binding regime; that is, the concentrations of immobilized Cra-DNA were near the K_d_ for FruK binding – 2.0 ± 0.5 x 10^−9^ M.

**Figure 4.**
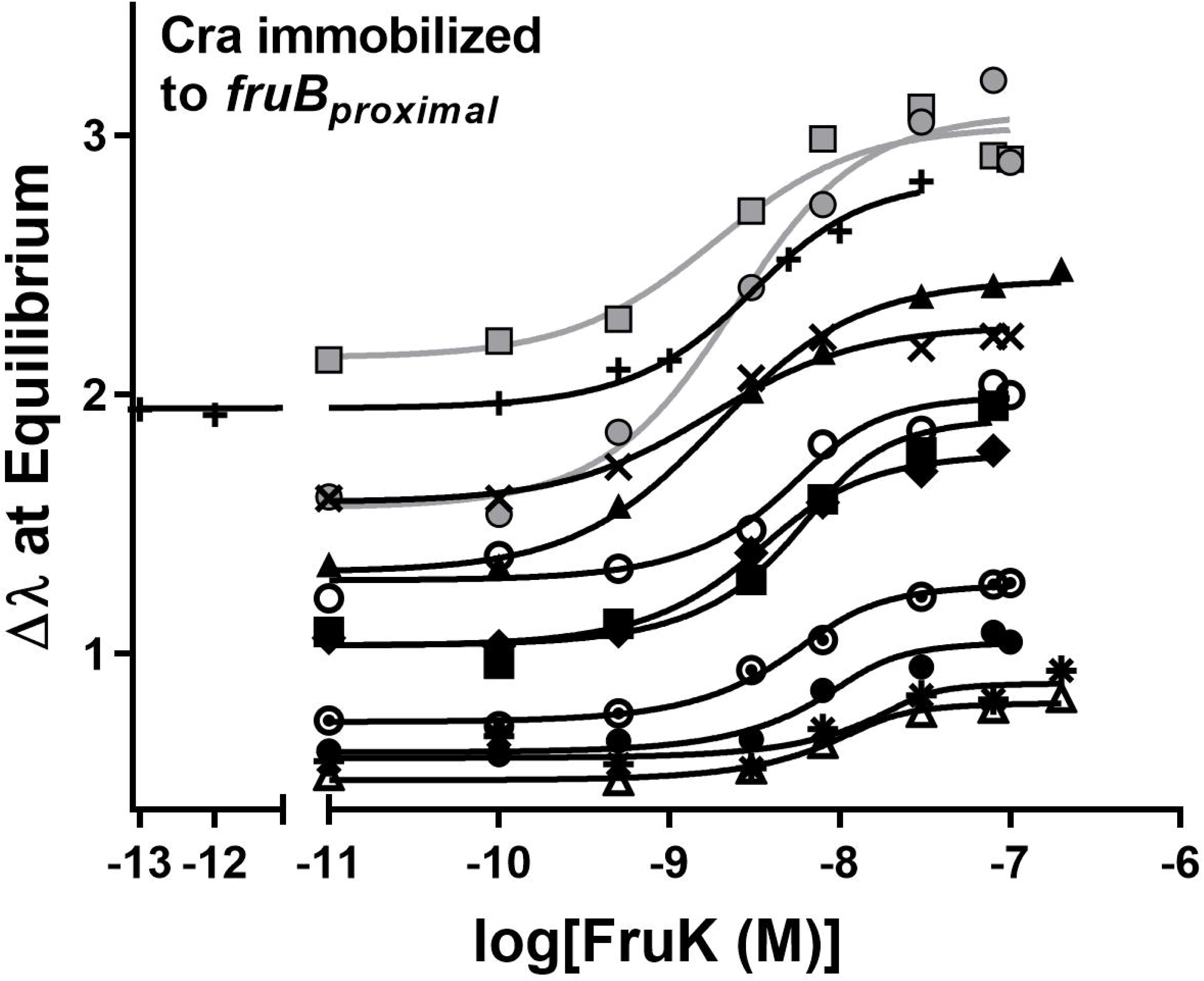
Purified FruK binds to immobilized, purified Cra/DNA. In these biolayer interferometry experiments, biotinylated DNA was immobilized to streptavidin coated fiber optic tips and then incubated with a constant concentration of Cra (2.5 nM) and varied concentrations of FruK (x axis). The individual binding curves denoted with the varied symbols are from independent experiments that used different lots of purified Cra and FruK. Curves were generated by global fitting of these independent assays to Equation 2: K_d_ was common to all curves, whereas all other parameters were unique to each curve (*i.e.*, each curve had its own value of DT, Y_max_, etc.). The variation in the Y axis intercept values arises from varied amounts of immobilized DNA; this variation is required in order to mathematically determine K_d_^25^ and either arose inherently due to differences among fiber optic tips (discussed extensively in ^25^) or was deliberately varied by diluting biotinylated-DNA with free biotin prior to immobilization. The global fit for 10 assays with untagged proteins (black symbols/fit lines) yielded a K_d_ of 2.0 ± 0.5 x 10^−9^ M. Including data for the His-FruK (gray circles and line) in the global fit yielded a K_d_ of 2.1 ± 0.4 x 10^−9^ M; a similar result was obtained for His-Cra (1.6 ± 0.4 x 10^−9^ M; gray squares and line). Experiments were performed in 50 mM HEPES, 100 mM KCl, 0.1 mM EDTA, 10 mM MgCl_2_, 0.3 mM EDTA, pH 7.

Since both wild-type Cra and the LLhF chimera interact with FruK with high affinity, we conclude that FruK indeed interacts with the regulatory domain(s) of Cra. Three control experiments indicate that this interaction is specific. (i) FruK alone did not bind to *fruB* DNA directly (Supplemental Figure 4, magenta). Neither Cra nor FruK bound to unrelated proteins, which rules out either protein being “sticky” and having promiscuous binding: (ii) As noted above, FruK did not bind full-length LacI^[25]^; (iii) DNA-bound Cra did not interact with *E. coli* pyruvate kinase, which is another metabolic kinase that is transcriptionally regulated by Cra^29^ (Supplemental Figure 4).

Finally, the affinity of FruK for wild-type Cra was nearly 10-fold tighter than that determined for LLhF^25^. Since FruK must bind regulatory domain of these two proteins, the altered binding affinity could arise from (i) the different the DNA binding domains of Cra and LLhF, which would be a long-range effect on FruK binding, and/or (ii) the different DNA operator sequences that were bound to Cra and LLhF, which would be an allosteric effect on FruK binding.

Next, we attempted to quantify FruK binding to Cra in the absence of DNA by directly immobilizing either His-tagged Cra or His-tagged FruK to BLI tips. However, neither protein could efficiently bind the untagged partner; equilibrium was not reached after several hours. The impeded binding could be due to either the His-tag or immobilization on the tips. Since the tagged proteins functioned like non-tagged protein in other versions of the binding assay (*e.g.,* Figure 4, gray symbols) and in cell lysate (Figure 3BC), we conclude that direct immobilization of either Cra or FruK onto the BLI tip imposed steric constraints that impaired binding.

### FruK dimer binds to a Cra dimer

To assess the stoichiometry of the Cra-FruK complex in the presence and absence of DNA, we used atomic force microscopy (AFM). This technique uses a cantilevered tip to measures the dimensions of individual molecules and complexes that have been deposited on a mica surface. Molecular volumes are then used to estimate molecular weights (MW) for each species. Prior to these experiments, the oligomerization state of unbound FruK was not known. Since its homolog ribokinase is a homodimer^30^, we hypothesized that FruK would also be a dimer. The minimal functional unit of Cra is a homodimer, which has the capacity to bind one DNA operator (*e.g.,* ^31–33^). The DNA for these experiments comprised a region of the *fruBKA* promoter that contains both the *fruB_distal_* and *fruB_proximal_* Cra operator sites; as such, one DNA had the potential to bind two Cra dimers.

Images of representative complexes are shown in Figure 5A and the complete AFM images are available at GitHub (https://github.com/ptoneil/2022_LSK_AFM_CraFruK.git). When the measured molecular volumes for the protein particles observed in the FruK+DNA and Cra+DNA samples were transformed into molecular weights, results corresponded very closely to the molecular weights expected for each homodimer (Figure 5B); this provided confidence in the assumptions used for computations^34–36^. The homodimeric Cra observed here differs from a previous report, in which purified Cra with a C-terminal His-tag appeared to be tetrameric^31^; perhaps these differences arise from different purification conditions.

**Figure 5.**
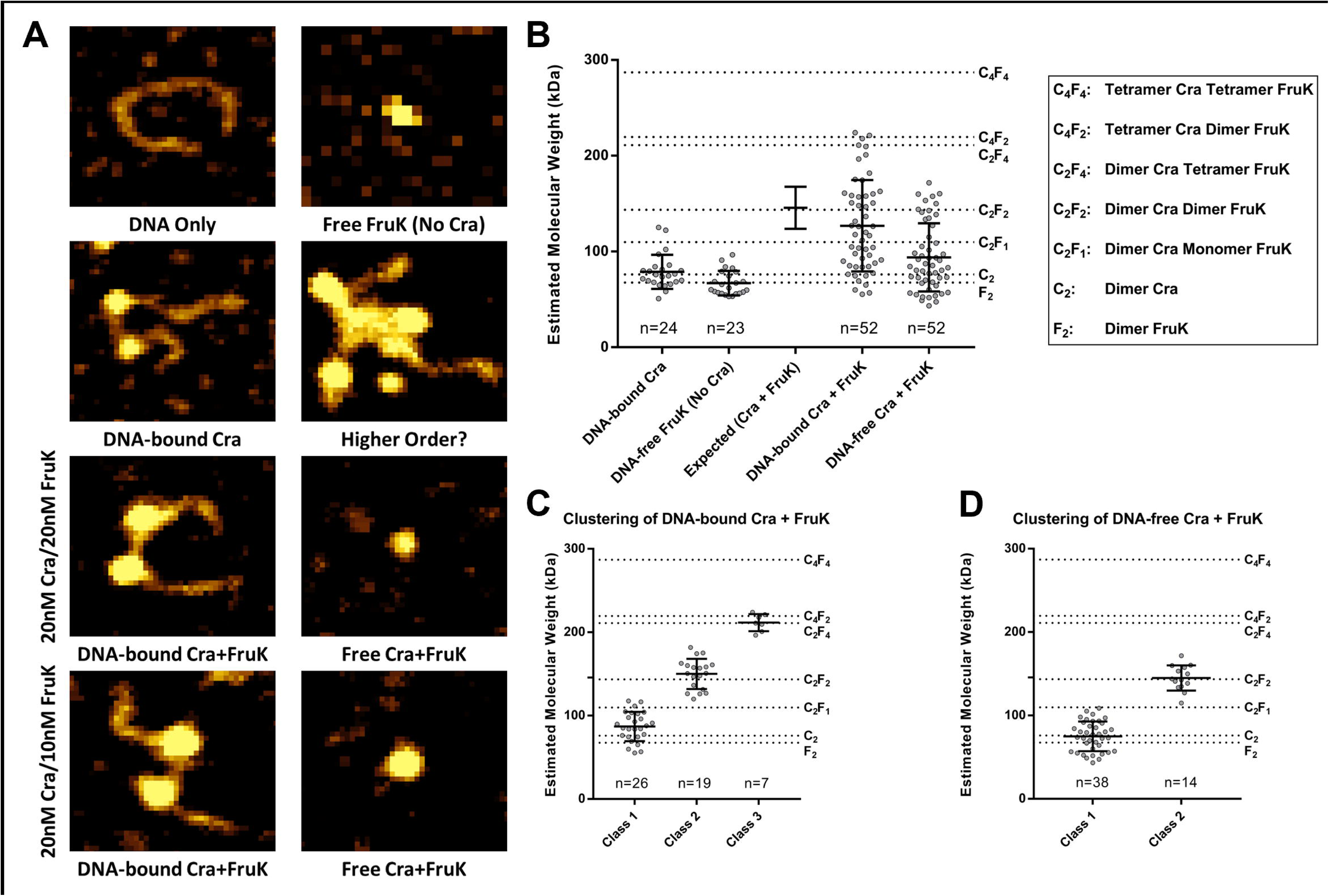
Distributions of Cra, FruK, and DNA complexes observed with atomic force microscopy. (A) Various combinations of DNA, Cra, and FruK were deposited on mica chips and scanned with AFM. The representative images shown represent 80 x 70 nm^2^ sections of the scanned mica; complete images can be found online (https://github.com/ptoneil/2022_LSK_AFM_CraFruK.git). (B) Size distributions of observed protein complexes. Each dot on this graph represents a unique, measured peak in an AFM image; the bar represents the mean and standard deviation for each population of dots. The number of individual complexes measured (n) is given above the x-axis. The dotted reference lines represent the expected molecular weights for various complexes. As another reference, values for the experimentally measured “DNA-bound Cra” and “Free FruK (No Cra)” were used to generate an “Expected (Cra + FruK)” value with propagated standard deviation. The sample containing DNA, Cra, and FruK were combined from two independent samples, one with a Cra:FruK ratio of 2:2 and the other with a 2:1 ratio. Separate analyses of these two samples are in Supplemental Figure 6. (C) The “DNA-bound Cra + FruK” population was well-fit using one-dimensional cluster analysis with a three-class system, fixing the upper class to n=7. Other fitting attempts are descripted in Supplemental Figure 7. (D) The “DNA-free Cra + FruK” population was well-fit using one-dimensional cluster analysis with two-classes.

Consistent with BLI results, FruK alone showed no binding to DNA (Supplemental Figure 5). As expected, Cra clearly bound to DNA (Figure 5A), often causing a “kink” in the DNA that matched the 45-50° bend angle that occurs when LacI/GalR proteins bind to their cognate operator sites (*e.g.,* ^37,38^). Several instances were observed in which both operators on a single DNA were bound by separate Cra homodimers (*e.g.,* Figure 5A).

When all three components (Cra, FruK, and DNA) were present in an AFM sample, the volumes of complexes ranged from that of (i) a single protein dimer “(dimer)_1_”, which could be either the Cra or FruK homodimer, (ii) to a dimer of protein dimers, “(dimer)_2_”, (iii) to a trimer of protein dimers, “(dimer)_3_” (Figure 5B). Model-free, one-dimensional clustering of these distributions identified (i) three distinct subpopulations of DNA-bound complexes, centered around (dimer)_1_ (presumably Cra), (dimer)_2_, or (dimer)_3_ (Figure 5C) and (ii) two-populations of DNA-free complexes, centered around (dimer)_1_ (which could be either Cra or FruK) or (dimer)_2_ (Figure 5D). Due to the similar molecular weights of the Cra and FruK homodimers, as well as their overlapping distributions of measured volumes, the composition of (dimer)_2_ and (dimer)_3_ cannot be directly elucidated. However, we assumed that every DNA-bound complex must contain at least 1 Cra homodimer. Furthermore, since only homodimers were observed in the single protein samples, we deduced that the (dimer)_2_ complex must contain one each of the Cra and FruK homodimers and that the (dimer)_3_ complex must contain a mixture of Cra and FruK dimers.

Additional information about the possible stoichiometry of (dimer)_3_ can be deduced from samples with varied protein concentrations (Supplemental Figure 6): The sample comprising 20 nM Cra/10 nM FruK had primarily (dimer)_2_ and some (dimer)_3_. When the FruK concentration was increased (20 nM Cra/20nM FruK), the population of particles shifted to *smaller* complexes. Thus, we deduce that (dimer)_3_ is Cra-FruK-Cra (each protein being a dimer), and that additional FruK saturated the Cra binding sites limiting complex formation to (dimer)_2_ Cra-FruK (Supplemental Figure 6).

Samples containing all three components (Cra, FruK, and DNA) also showed even larger complexes that were mostly absent from the Cra+DNA and FruK+DNA samples (*e.g.,* Figure 6; other images available on GitHub). These were excluded from quantitative analyses based upon their size and complexity. While these larger complexes may represent non-specific aggregation of Cra and FruK, the fact that all samples were made within minutes of each other and the single protein samples did not contain these complexes, makes this unlikely. Rather, the presence of higher order oligomers raises the possibility that a FruK dimer may bridge Cra dimers (*i.e.*, (dimer)_3_) to enable “looped” complexes with two or more DNA operators, similar to the dual-operator loops known to enhance repression by tetrameric LacI (*e.g.*, ^26,39–46^). Indeed, several of the Cra/FruK aggregates appear to include at least two DNA operators.

**Figure 6.**
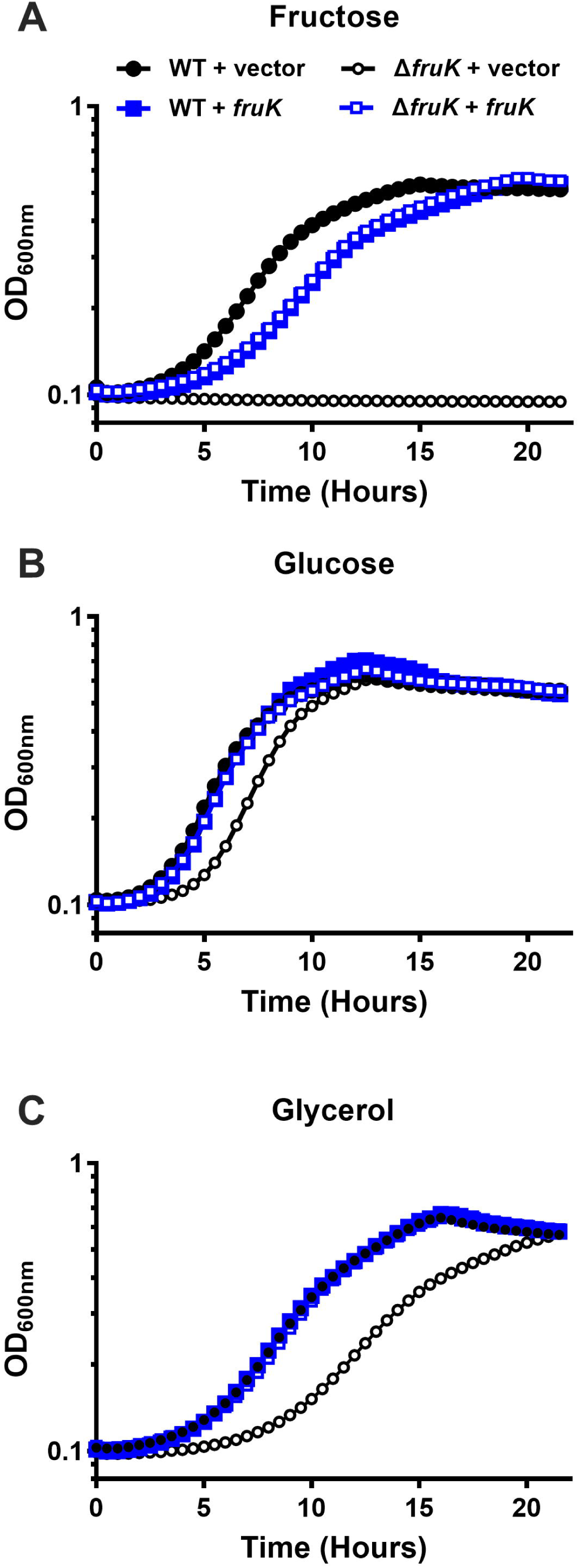
FruK contributes to *E. coli* metabolism. Bacterial growth was measured for wild-type and Δ*fruK724::kanR* (Δ*fruK*) strains of *E. coli* transformed with pHG165a parent plasmid (vector) or pHG165a-FruK that constitutively expressed FruK (*fruK*). Cultures were grown in M9 minimal media containing either 0.2% fructose (A), glucose (B), or glycerol (C). OD_600_ was measured at 30-minute intervals for 22 hours. Symbols represent the mean with standard deviation for n=3 replicates; error bars are smaller than the symbols. The data shown are representative of three independent experiments, which were in good agreement with each other: For the uncomplemented Δ*fruK* strain relative to wild type in media with glucose, the time required to reach half maximal OD_600nm_ was delayed by 0.5, 1.3, 1.7 hours, respectively; for media with glycerol, the time required to reach half maximal OD_600nm_ was delayed by 3.3, 3.4, and 3.8 hours, respectively. Constitutive over-expression of FruK from a plasmid greatly reduced or abolished the delay time in all three experiments. Results in different growth conditions (Supplemental Figure 8) are in good agreement with the results shown here.

### The absence of FruK impacts growth on a non-preferred carbon source

If either the reverse FruK reaction (F-1,6-BP to F-1-P) or the direct interaction of FruK with Cra are biologically relevant, deleting FruK should have biological consequences when *E. coli* are grown on non-fructose carbon sources. Based on its potential to generate Cra’s allosteric regulator F-1-P from the central metabolite F-1,6-BP, we hypothesized that abolishing expression of FruK protein would alter Cra’s ability to regulate switching among the metabolic pathways that enable efficient use of alternative carbon sources. Indeed, an early report for a *fruK* mutant strain showed growth defects on lactate and a corresponding decrease in PEP-synthase (an enzyme required for gluconeogenesis^47^ whose transcription is activated by Cra^48^).

For our growth experiments, we used the Δ*fruK724*::*kanR* strain from the Keio collection^49^. To test whether FruK impacts growth on alternative carbon sources, we sub-cultured strains from M9/glucose media to M9 media with either glucose, glycerol, or fructose as the sole carbon source. In previous reports, ablation of fructose-1-kinase activity abolished *E. coli* growth on fructose (*e.g.,* ^11^); thus, as a control experiment, we confirmed that the wild type, but not the Δ*fruK724*::*kanR* mutant strain, was able to grow on fructose (Figure 6A black). Growth on fructose was restored when FruK was constitutively expressed from a plasmid. Surprisingly, FruK expression on the multi-copy plasmid slightly inhibited growth on fructose in both the wild-type and Δ*fruK724*::*kanR* strains (Figure 6A, open and filled blue symbols) relative to wild-type strain.

Next, we examined the ability of the strains to grow on the preferred carbon source, glucose (Figure 6B). In this case, the Δ*fruK724*::*kanR* strain showed a slightly longer lag phase (Figure 6B, open black symbols) relative to the other strains. This lag phenotype was reverted when FruK was provided on a plasmid (Figure 6B, open blue symbols). Finally, we examined how the absence of FruK impacts growth on the non-preferred carbon source glycerol (Figure 6C). Under this condition, the Δ*fruK724*::*kanR* strain showed a pronounced lag phase (and somewhat slower growth rate) than the other samples (Figure 6C, open black symbols); complementation with wild-type FruK again restored wild-type growth kinetics.

Thus, these results show that deleting FruK alters growth on glucose and glycerol, and this phenotype is exacerbated on the non-preferred carbon source. In separate experiments, we examined the transition from LB to minimal media with different carbon sources for wild type and the Δ*fruK724*::*kanR* strains. Similar to the other experiments, the absence of FruK led to a delayed in growth (Supplemental Figure 8), consistent with FruK playing a role in metabolic switching. These combined results are consistent with FruK playing a role in central metabolism that is beyond that of fructose metabolism.

### Biofilm formation is altered in the ΔfruK724::kanR strain

Another phenotype that requires precise regulation of central metabolism in *E. coli* is biofilm formation.^50–52^ In addition to controlling the switch between glycolysis and gluconeogenesis that occurs during biofilm formation, Cra directly activates the expression of the *csgDEFG* genes that are necessary to produce Curli, an attachment filament that is important for biofilm formation^53^. Indeed, a *cra* mutant strain is deficient in biofilm formation.^53^ If FruK only acted in fructose metabolism, its knock-out would *not* be expected to alter biofilm formation (unless fructose was included in the growth media). However, if FruK plays a larger role in regulating central metabolism, its knock-out might alter biofilm formation.

Thus, we next tested whether the absence of FruK would impact biofilm formation. Indeed, the Δ*fruK724*::*kanR* strain had decreased biofilm formation when grown in the presence of glucose (Figure 7 white). No difference in biofilm formation was observed between the wild-type and Δ*fruK724::kanR* strains when grown in media with glycerol as a carbon source (Figure 7 gray). These results are consistent with the absence of FruK altering Cra’s activity and further demonstrates an impact on cell processes beyond FruK’s role in fructose metabolism, most likely through its effect on Cra.

**Figure 7.**
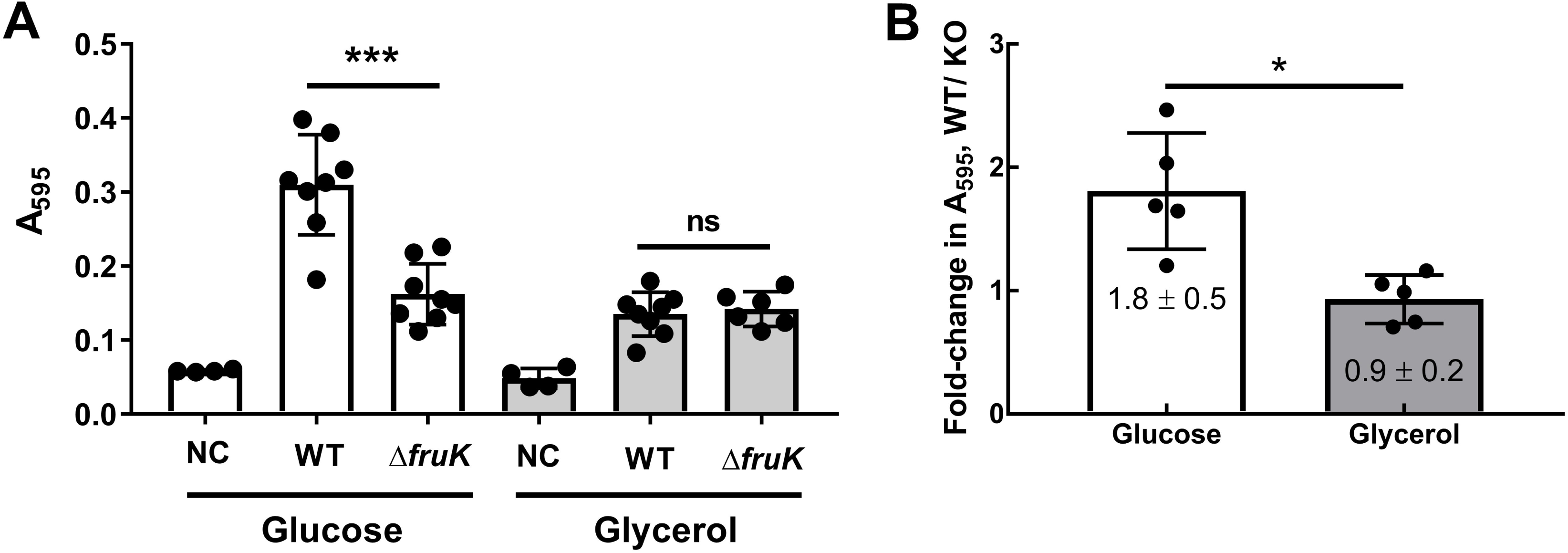
FruK contributes to biofilm formation. Biofilm assays with uncomplemented wild-type (WT) and Δ*fruK724::kanR* (Δ*fruK*) strains of *E. coli* were performed in M9 media with the indicated carbon source after incubation at 37°C for 72 hours. “NC”: no bacterial cells inoculated. (A) In this representative experiment, each symbol represents a technical replicate (4 to 8) and the column represents the mean with SD. “***” indicates a significant difference (p=0.0002) between strains by Welch’s t-test. (B) The average fold-change in A_595_ between WT and Δ*fruK* strains grown on glucose or glycerol for five independent experiments are plotted as bars. Each dot represents the average of multiple technical replicates (*e.g.,* the average from panel (A)); the error bars associated with each independent experiment (dot) are not shown here but were propagated to calculate the standard deviation of the full dataset that is shown with error bars. “*” indicates a significant difference (p=0.01) between growth conditions by Welch’s t-test.

## Discussion

Although studied for decades, our understanding of how central metabolism is regulated in *E. coli* remains incomplete. Here, we considered that the little-studied enzyme FruK might play additional roles beyond that of fructose catabolism. Indeed, results for the growth assays showed that wild-type FruK plays a broader metabolic role, perhaps even contributing to metabolic switching that is known to the regulated by Cra. A second phenotype arising from disrupted FruK expression was altered biofilm formation. These phenotypes might be accomplished by disrupting any (or several) of five different molecular mechanisms:

First, the reverse FruK reaction demonstrated herein (Figure 2) generates the strong Cra allosteric inhibitor F-1-P from physiologically relevant concentrations of F-1,6-BP, thereby providing a metabolic pathway by which other sugars can be used to create F-1-P. The reverse reaction does not appear to be influenced by interaction with Cra, suggesting that the FruK region implicated in Cra binding does not impact FruK catalysis.

Second, the nanomolar interaction observed between FruK and DNA-bound Cra is tight enough to exist *in vivo*. Cra and FruK have long been known to co-regulate each other’s activities *via* transcription repression and F-1-P production, respectively (Figure 1). The current results raise the intriguing possibility that the direct Cra-FruK interaction could also play an important biological role. If nothing else, the direct Cra-FruK interaction would produce F-1-P in the immediate vicinity of Cra. In addition, FruK binding to Cra might alter its ability to regulate transcription for specific operons. Since Cra directly regulates >40 natural operators, large scale studies will be needed to test this hypothesis. One means by which this could occur is through altering the binding affinity between Cra and a subset of its target operators.

Indeed, a third possibility is that the higher order Cra-FruK-DNA complexes observed in AFM could simultaneously bind and loop multiple DNA operators, similar to LacI’s regulation of the *lac* operon (*e.g.*, ^26,39–44^). Consistent with this, six promoters in the Cra regulon have multiple Cra binding sites^4,5,7,53–55^. To test this hypothesis, additional experiments will require assays with longer inter-operator spacing than the current construct^56,57^.

A fourth possibility is that the Cra-FruK interaction could alter the Cra interaction with other transcription factors. Operon-specific interactions are known for enterohemorrhagic *E. coli* Cra and KdpE^58^, as well as for *E. coli* Cra and FNR^59^. Fifth, for various Cra-activated pathways (such as those by which glycerol and lactate are metabolized), Cra-FruK binding could facilitate interactions with some other factor, such as RNA polymerase, as observed for members of the AraC/XylS family of transcription activators (*e.g.*, ^60–62^).

Finally, we anticipate that FruK’s newly discovered role(s) will vary with changing metabolic conditions. As noted above, *fruK* gene expression is itself regulated by Cra, leading to different cellular concentrations of FruK under different metabolic conditions. Such changes could alter whether the direct Cra-FruK complex is formed; indeed, this was likely detected in the *in vivo* pull-down assay first used to detect the Cra-FruK interaction (Figure 3A; Supplemental Results). F-1,6-BP concentrations also depend upon available carbon sources^22,27^ (*e.g.,* Supplemental Figure 1 and Supplemental Table 1). Effects of F-1,6-BP concentration changes on the reverse FruK reaction might resolve the conundrum as to how F-1,6-BP signals the need for altered metabolic flux. Thus, once the molecular mechanism(s) of FruK’s contributions are elucidated, results should be used to improve current models of *E. coli* metabolic flux^16,21,22^ to determine whether discrepancies with measured metabolites have been resolved.

In summary, the intriguing range of ways that FruK might co-regulate *E. coli*’s central metabolism will be important to further consider. As both Cra and FruK are conserved within the clade of γ-proteobacteria (*e.g.*, ^8^), co-regulation by FruK could play a key role in a wide range of organisms.

## Materials and Methods

### Common Reagents

The trisodium salt of D-fructose-1,6-bisphosphate (F-1,6-BP), ATP, and ADP were purchased from Sigma-Aldrich (St. Louis, MO). For experiments, F-1,6-BP, ATP and ADP nucleotides were dissolved in the appropriate buffer (detailed below) and the pH was adjusted to 7.0.

The barium salt of F-1-P was purchased from Sigma-Aldrich (St. Louis, MO). To generate the more soluble sodium salt, Ba^+^ F-1-P was subjected to ion exchange chromatography to using a Dowex 50WX4 (Sigma) column pre-charged with 2 N HCl and equilibrated in ddH_2_O. After several passages, the pH of the final eluant was adjusted to 7 with NaOH. Eluted F-1-P was lyophilized and used to make saturated stock solutions in ddH_2_O. Concentrations of the stock solutions were determined by stoichiometric implementations of the enzyme assay, as reported previously^25^.

The various operator sequences used are shown in Table 1. For pull-down, BLI, and filter binding assays, DNA oligos were purchased from Integrated DNA Technologies (Skokie, Illinois). Single stranded DNA was annealed using the protocol described in Swint-Kruse *et al.*^63^. For pull-down assays and biolayer interferometry, the purchased DNA oligos were 5’ biotinylated. For filter binding assays, annealed DNA was labeled with 32P-ATP (*e.g.*, ^63^). For atomic force microscopy, a 331 bp of *fruBKA* promoter, spanning both the fruB_distal_ and fruB_proximal_ binding sites, was amplified from the DH5α (F^−^ φ80*lac*ZΔM15 Δ(*lac*ZYA-*arg*F)U169 *rec*A1 *end*A1 *hsd*R17(r_K_ ^−^, m_K_ ^+^) *pho*A *sup*E44 λ^−^*thi*-1 *gyr*A96 *rel*A1) strain of *E. coli*. Further details of this reaction are described in the Supplemental Methods: Preparation of DNA for AFM.

**Table 1.**
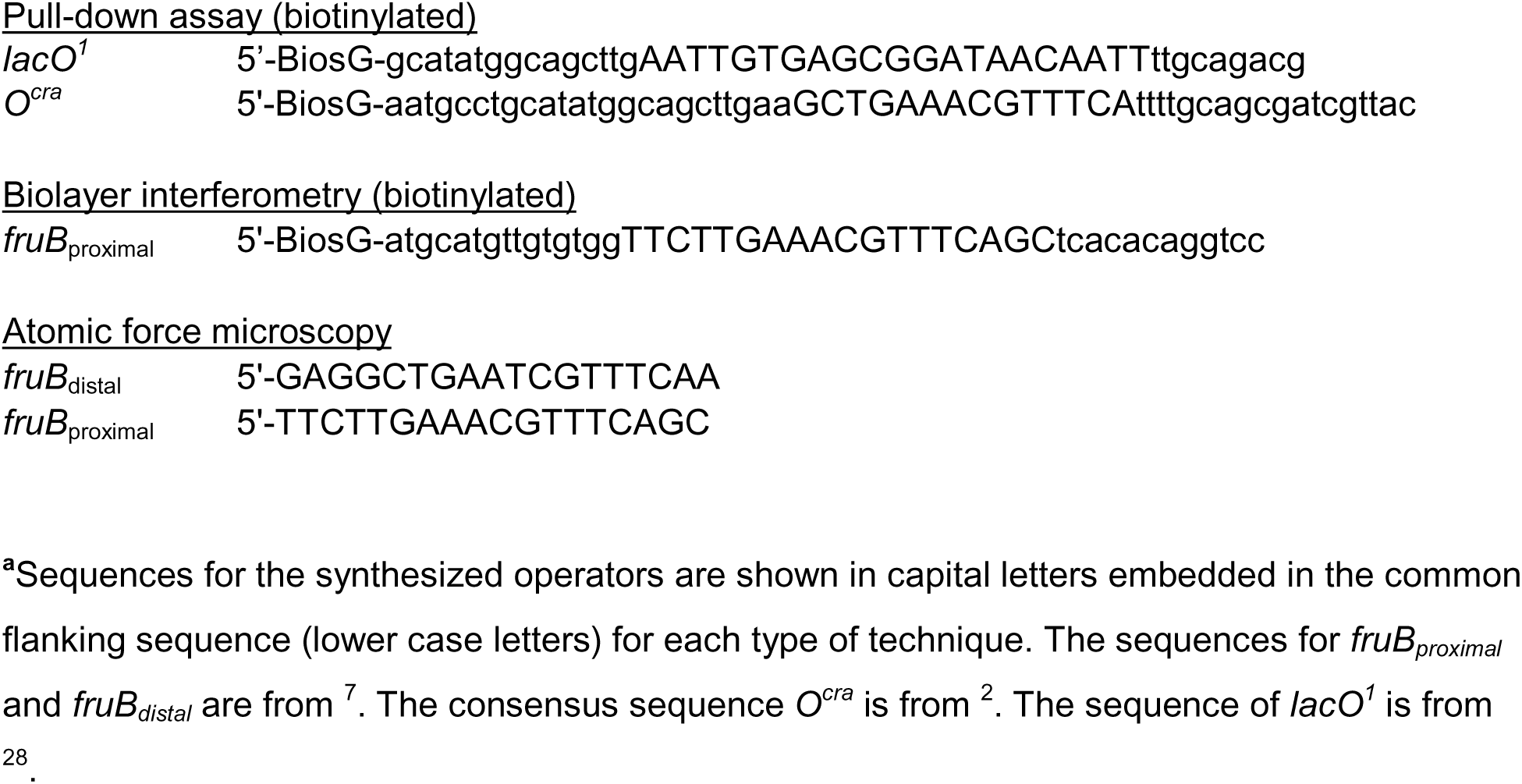
Operator sequences used in Cra/FruK experiments.^a^.

### Plasmid construction

Lists of bacterial strains and plasmids used in this work are in Supplemental Table 2. The coding region for wild-type Cra (Supplemental Sequences) was amplified from Hfr(PO1), *lacI22*, λ-, *e14-, relA1, spoT1, thiE1 E. coli* (strain name 3.300; *E. coli* Genetic Stock Center, Yale University) using the primers 5’-ggtctccaggtgtgaaactggatgaaatcgc and 5’-gagctcttagctacggctgagcacgccgcgg. The gene was subcloned into the pHG165a plasmid using the procedure of Jones^64^. Full gene sequencing showed that the coding region exactly matched that of UniProt P0ACP1. This plasmid has been deposited with Addgene (90066; plasmid name FruR-pHG165a).

An expression plasmid carrying the *E. coli fruK* gene on pET-3a was a gift from the Van Schaftingen laboratory at the Université Catholique de Louvain, Belgium^24^ (Addgene ID 186256). The FruK amino acid sequence was determined for this plasmid and is reported in Supplemental Sequences. Sequencing was performed by ACGT, Inc (Wheeling, IL).

To generate pET3a-Cra-C-His, pET3a-N-His-FruK, and pET3a-FruK-C-His for BLI trials, plasmids expressing coding regions for untagged Cra and FruK were used as templates to amplify their respective genes, using 2 overlapping primers that include repetitive histidine codons (Supplemental Table 3). The amplified products were digested with DpnI (New England Biolabs) to remove the parental plasmid before transformation into DH5-alpha *E. coli*. Positive clones were validated by gene sequencing. pET28a versions of enterohemorrhagic *E. coli* (EHEC) FruK*-*C-His and EHEC Cra-C-Flag were generated by similar methods.

The open reading frame encoding *E. coli* FruK was amplified from EHEC and cloned into pHG165a for constitutive, low-level expression^26^ and subcloned into pET28a plasmid for inducible expression using the ABC cloning method^65^. His-tagged versions were also created for pull-down assays. Primers used to generate pHG165a*-*FruK, pHG165a*-*FruK-C-His, and pET28a-FruK-C-His are listed in Supplemental Table 3.

### Analyses of reported FruK sequences

Upon sequencing the pET3a FruK expression plasmid, we found the coding region contained differences from the reported UniProt sequence (P0AEW9) (Supplemental Sequences). To assess whether the pET3a (or any other) FruK variant sequences occur naturally, we used the pET3a FruK sequence as the query in a BLAST search^66^ of the nr database. The pET3a FruK sequence has 100% identity with each of the three sequences reported for *Salmonella* FruK. In addition, the sequences reported for *E. coli* FruK showed a range of mutations, including those observed in the pET3a/*Salmonella* sequence. To survey this range, BLAST results were filtered between 80-99.9 percent identity with query coverage between 85-100% of the FruK query. The resulting multiple sequence alignment was edited for aberrant sequences (generally comprising sequences with long N-termini or inserts of >20 amino acids). The remaining 364 sequences were aligned using clustalOmega^67^ and analyzed with ConSurf^68^ to assess the variability at each FruK position; 212 of the 312 positions showed some variability; only 100 positions were absolutely conserved. Several representative sequences are shown in Supplemental Sequences, including the FruK sequence on pET3a and that cloned from EHEC to pHG165 and pET28a. While sequencing errors are always possible, the frequency of observed changes leads us to conclude that *E. coli* FruK experiences natural variation.

### Purification of FruK and Cra; verification of their activities

The procedure for purifying FruK was adapted from^24^, as reported previously in the Supplementary Material to Weeramange *et al.*^25^. In brief, 1 L cultures of BL21 (DE3) pLysS *E. coli* [F–, *omp*T, *hsd*S_B_ (r_B_–, m_B_–), *dcm*, *gal*, λ(DE3), pLysS, Cm^R^, [malB^+^]_K-12_(λ^S^)] carrying the FruK-pET-3a expression plasmid were grown to an OD_600_ of ~0.6, at which time they were induced with 1 mM ITPG and grown for 44-45 hours at 22°C. After lysis in breaking buffer (20 mM potassium phosphate, 5 mM EDTA, 1 mg mL^−1^ lysozyme, 1 Pierce Protease Inhibitor Tablet (Thermo Scientific, Rockford, IL), 1 mM DTT, pH 7.4), FruK protein was purified with a two-step ammonium sulfate precipitation (using the supernatant from a 30% precipitation and the pellet from a 70% precipitation). The pellet was resuspended in Buffer A (20 mM HEPES, 20 mM KCl, ½ tablet Pierce Protease Inhibitor A32963 from Thermo Scientific, 10% glycerol) and desalted with either (i) a Sephadex G25 column or (ii) by diluting the ammonium sulfate resuspension 1:5 with 20 mM KCl Buffer A and then reducing the supernatant volume by half, using a Vivaspin MWCO 10 kDa concentrator (Sartorius Stedim Lab Ltd, Stonehouse, UK) and centrifugation with a Sorvall HB-6 rotor (Thermo Fisher) at 5500 rpm (4900xg) for 12-15 minutes at 4°C. The latter performed better in the following subsequent DEAE ion exchange chromatography. For anion exchange, a two-step gradient was used to elute the FruK: (i) 20-250 mM KCl in Buffer A and (ii) 250-500 mM KCl in Buffer A. FruK protein elutes near the end of the first gradient. The identity of the purified FruK protein was confirmed by mass spectrometry by the KUMC Mass Spectrometer/Proteomic Core.

Aliquots of eluted FruK were stored in 30% glycerol at −80°C. Glycerol is a key buffer component. If this osmolyte was removed by dialysis, FruK enzymatic activity decayed over the course of 30 min – 1 hour. Prior assays indicated that part of the FruK elution peak was contaminated with a small molecule that inhibits various FruK functions. Thus, immediately prior to each assay, this contaminant was removed by diluting the protein to ≤ 3.5×10^−3^ mg mL^−1^ (~100 nM) with FruK exchange buffer (20 mM HEPES, 40 mM KCl, 1 mM EDTA, 20% glycerol, 0.3 mM DTT) and re-concentrating it with a VivaSpin MWCO 10 kDa; dilution and concentration was carried out 3 times in total. The final protein concentration was determined with a Bradford assay (Bio-Rad Laboratories, Inc, Hecules, CA) and is reported as molar dimer. The enzymatic activity of the purified protein was verified using the reaction designed for the forward enzymatic reaction, as previously reported^25^.

The protocol to purify wild-type Cra was similar to that used to purify lactose repressor protein (LacI) and LLhF^25,63,69–72^. *E.coli* B *omp*T, *hsdS_B_*(*r_B_*^−^*m_B_*^−^)^−^, *gal, dcm, lac* cells (strain name “BLIM”^73^ were transformed with plasmid constitutively expressing wild-type Cra and grown in 6 L 2xYT media for 19 hours at 37°C; for some preps, growth media included 1% fructose, but this did not appear to alter either the amount or activity of the purified Cra. After pelleting by centrifugation at 676xg for 20 minutes and 4°C, the cells were resuspended in ~50 mL breaking buffer (0.2 M Tris-HCl, 0.2 M KCl, 0.01 M MgCl_2_, 1 mM DTT, 5% glucose, pH 7.6) with 20 mg lysozyme and half of a crushed Pierce Protease Inhibitor Tab (Thermo Scientific, Rockford, IL) and frozen at −20°C. All subsequent purification steps were carried out on ice or at 4°C.

Cell lysis was accomplished by thawing the cell pellet on ice with breaking buffer added to a final volume of 150 mL and another crushed protease tab. At this step, the addition of 120 mg extra lysozyme was critical for chaperoning the Cra through ammonium sulfate precipitation, which was an unexpected activity of lysozyme. After lysis, 240 μL of 80 mg mL^−1^ DNase I from bovine pancreas (Roche Diagnostics, Mannheim, Germany) was added along with MgCl_2_ to a final concentration of 30 mM. The crude cell extract was cleared by centrifugation at 9700xg (8,000 rpm) for 50 min at 4°C prior to 23% ammonium sulfate precipitation (Fisher Scientific, Fair Lawn, NJ). The technical aspects of ammonium sulfate precipitation were very important. The chemical must be finely ground and lightly salted over the whole surface of the solution (generally 180-200 mL in a 400 mL beaker) at a rate so slow that crystals never reach the bottom of the flask. The mixture was allowed to precipitate for 30 minutes at 4°C with slow stirring prior to centrifugation at 9700xg (8,000 rpm) for 40 min and 4°C.

After precipitation, the pellet was gently resuspended in 50 mL cold 0.05 M KP buffer (KP buffer comprises potassium phosphate at the indicated molarity, 5% glucose, 0.3 mM DTT, pH 7.5) and dialyzed in regenerated cellulose Fisherbrand Dialysis Tubing (MWCO 12,000-14,000) against 1 L of cold 0.05 M KP buffer for 30 minutes, with three buffer changes. The dialysate was cleared by centrifugation, diluted 1:6 with 0.05 M KP buffer, and loaded onto a phosphocellulose column (Whatman P11) that was pre-equilibrated in 0.05 M KP buffer. The Cra protein eluted shortly after the beginning of a 0.05 M to 0.2 M KP buffer gradient, which was benchmarked against LacI variants as behaving like a dimer^63,74^. Purity was assessed on SDS-PAGE with silver stain and was estimated to be >90% (*e.g.*, Supplementary Figure 2 in ^75^). The pre-column dilution (1:6 in 0.05 M KP buffer) was key to good purity; without it, Cra eluted from the phosphocellulose column much later and was contaminated with lysozyme and other proteins. Purified Cra was aliquoted and stored at −80°C. Protein concentration was determined by Bradford assays (Bio-Rad Laboratories, Inc. Hercules, CA) and was generally 0.1-0.2 mg mL^−1^. DNA activity assays^76^ using the filter binding technique (below) were used to assess the fraction of purified Cra that was active; preparations were generally 70-90% active.

Note that the concentrations of purified FruK and Cra are given in terms of molar dimer for all assays.

### FruK Reverse Reaction

The forward reaction of purified FruK was detected by coupling ADP formation to the activities of pyruvate kinase and lactate dehydrogenase, ultimately assessing the change in NADH concentration (*e.g.,* Supplemental Figure 2)^25^. To detect the formation of ATP during the reverse reaction, we used the Kinase-Glo kit (Promega, V6728) to assess the activity of purified FruK.

To maintain both good buffer capacity and a constant ionic strength across all substrate concentrations, stock solutions of trisodium F-1,6-BP (Sigma-F6803) and monosodium ADP (Sima-A2754) were made in Cra1X base buffer (50mM HEPES, 0.1 KCl, 0.1 EDTA, 10 mM MgCl_2_, pH 7). ATP (Sigma A9187) for controls were diluted in buffer containing Cra1X base buffer with 115 mM NaCl and 2 mM MgCl_2_. All substrate solutions were adjusted to pH 7.0. Assay dilutions were performed with Cra1X base buffer with 180 mM NaCl, 40 mM KCl and 10 mM MgCl_2_. The final ion concentrations in the assay wells were 66.6 mM K^+^, 95.5 mM Na^+^, 10 mM Mg^2+^, and 105 mM Cl^−^. The experiment shown in Figure 2 also included 275 nM “washed” Cra and 30 nM *fruB_distal_* DNA.

Triplicate technical replicates for each substrate concentration were prepared in Costar white 96 well plates as follows: Each well contained either (i) 60 μL of the FruK reaction sample and 50 μL Kinase-Glo reagent for a total volume of 110 μL or (ii) 60 μL of ATP at varying concentrations plus 50 μL of Kinase-Glo reagent for a total of 110 μL. The final FruK and Cra concentrations were maintained at 275 nM, ADP was 5.5 mM, F-1,6-BP ranged from 0.1 to 15 mM, and ATP controls ranged from 0.03 to 4 mM. For reactions that included Cra or DNA, these components were included in the dilution cocktail used for serial dilutions of F-1,6-BP. After incubating for 1 hour (shorter times produced lower signals), luminescence was read at room temperature using a SpectraMax M5 plate reader at all wavelengths.

Control reactions showed that that increasing the ionic strength inhibited activity of luciferase, making it critical to balance the counter ions of the changing substrate concentration (Supplemental Figure 9). Luciferase assays of the FruK forward reaction to monitor the disappearance of ATP did not detect enzyme activity. Since the forward reaction was readily detected in a different assay (Supplemental Figure 2), we concluded that either (i) the conditions needed to balance ionic strength across the titration of the forward reaction was even more inhibitory to the luciferase reaction and/or (ii) reagents in the luciferase reaction inhibited the FruK forward reaction.

### DNA pull-down assay with wild-type Cra and endogenous FruK

For pull-down assays, the pHG165a plasmid encoding wild-type Cra (Addgene ID 90066) was transformed into the *E. coli* strain 3.300 (genotype Hfr(PO1), *lacI22,* λ*-, e14-, relA1, spoT1, thiE1*). Cultures were grown at 37°C to saturation in 3 mL of MOPS media (Teknova, Hollister, CA: 40 mM morpholinopropanesulfonic acid, 10 mM NH_4_Cl, 4 mM tricine, 50 mM NaCl, and other trace metals listed for product number M2101) supplemented with 0.8% glycerol, 1.32 mM dibasic potassium phosphate, 10 mM NaHCO_3_, 0.2% casamino acids, 0.0025% thiamine and 100 μg mL^−1^ ampicillin^77,78^. Cells were pelleted by centrifugation, resuspended in 100 μL breaking buffer (0.2 M Tris-HCl, 0.2 M KCl, 0.01 M MgCl_2_, 1 mM dithiothreitol [DTT], 5% glucose, pH 7.6) and lysed via freeze-thaw with 40 μL of 5 mg mL^−1^ lysozyme. Crude cell lysates were clarified by centrifugation. Ten (10) μL supernatant were incubated with biotinylated DNA that was immobilized to streptavidin magnetic beads (1 pmol DNA to 1 μL beads; New England Biolabs, Ipswich, MA), as described in Meinhardt *et al.*^79^. Sequences of the biotinylated DNA operators (Integrated DNA technologies, Coralville, IA) are listed in Table 1. After washing in FB buffer (10 mM Tris HCl, 150 mM KCl, 10 mM EDTA, 5% DMSO, 0.3 mM DTT, pH 7.4), the protein-DNA-beads were resuspended in 15 μL of denaturing buffer (10 μL 20% sodium dodecyl sulfate plus 5 μL 1M dithiothreitol [DTT]). Adherent proteins were visualized with SDS-PAGE.

In prior pull-down assays with engineered LLhF, the second, slightly smaller band was identified with mass spectrometry to be FruK^25^. These experiments were carried out using the now-discontinued Phast-gel 10-15% gradient, (GE Healthcare, Piscataway, NJ; former catalogue #: 17-0540-01) to resolve the repressor protein band from the FruK band.

Results are in Figure 3A. The uncropped gels associated with this Figure are in Supplemental Figure 10.

### Pull-down assay with over-expressed FruK and Cra

BL21 (DE3) *E. coli* (Novagen) carrying a modified pET28a expressing either EHEC FruK-C-His or EHEC Cra-C-Flag were grown in 150 mL LB supplemented with kanamycin (50 µg mL^−1^) to an OD_600_ of 0.5. Protein over-expression was induced with isopropyl β-d- thiogalactopyranoside (0.5 mM) for 4 hours at 30°C. After centrifugation, the bacterial pellet was suspended in 4 mL of 50 mM sodium phosphate, pH 8.0, 0.5 mg mL^−1^ lysozyme and incubated on ice for 30 min. An equal volume of 50 mM sodium phosphate, pH 8.0, 2 M NaCl, 8 mM imidazole, 20% glycerol, and 0.5% Triton X-100 was added. After 30 min on ice, the bacterial lysate was sonicated then clarified by centrifugation.

For the pull-down assay, equal amounts (200 µl) of FruK and Cra cell lysates were mixed with 100 µl of nickel-nitrilotriacetic acid beads (Qiagen). After overnight incubation at 4°C with gentle rotation, mixtures were loaded on a Poly-Prep Chromatography Column (Bio-Rad) and washed with 5 mL of 50 mM sodium phosphate, pH 8.0, 600 mM NaCl, 10% glycerol, 60 mM imidazole. Adherent proteins were eluted in 50 mM sodium phosphate, pH 8.0, 600 mM NaCl, 10% glycerol, and 250 mM imidazole.

Elution samples were subjected to SDS-PAGE on a 15% acrylamide gel (Supplemental Methods: SDS-PAGE) stained with Coomassie Brilliant Blue R-250. These gels were made using the original method of Laemmli^80^. Aside from the Phast-gel used in the DNA-pull-down experiments (described above), no other commercial acrylamide gels tested to date have been able to resolve Cra from FruK due to their similar molecular weights. Western blot analysis used anti-His (Santa Cruz Biotechnology sc-8036; mouse) and anti-Flag (Sigma F7425; rabbit) as the primary monoclonal antibodies. Secondary antibodies were purchased from Li-Cor (goat anti-mouse IRDye 680RD and goat anti-rabbit IRDye 800CW). Western blots were imaged using a Li-Cor Odyssey at 680 and 800 nm.

Results are in Figure 3BC. The uncropped gels associated with this Figure are in Supplemental Figure 10.

### Biolayer Interferometry

In biolayer interferometry (BLI), macromolecules are immobilized to a fiber optic tip, which can then be dipped into solutions of other macromolecules. Changes to the thickness of the bound layer (*e.g.,* when a second protein binds the immobilized macromolecule) are detected by a change in reflectance when light is passed through the tip^81^. BLI was originally devised to measure the on and off rates of kinetic assays and thereby quantify binding affinities^82^; we previously adapted BLI to measure binding affinities under equilibrium conditions^25^. This adaption was required because the kinetic association curves for the Cra homologs LacI and LLhF were multiphasic^25^; furthermore, dissociation curves could reflect both dissociation of FruK from Cra and dissociation of Cra from DNA. The equilibrium implementation was used here to measure FruK binding to DNA-bound Cra, using an Octet RED96 (Forté Bio, Pall Life Sciences, Menlo Park, CA; Figure 4).

For FruK binding to DNA-bound Cra, experiments used “High-Precision Streptavidin” probe tips (Forte Bio) to which various biotinylated DNA sequences were attached (Table 1). Tips were first hydrated for at least 10 minutes in the DNA immobilization buffer (50 mM HEPES, 0.1 M KCl, 0.1 mM EDTA, 10 mM MgCl_2_, pH 7.0, 0.3 mM DTT with 0.1 mg mL^−1^ BSA), loaded with biotinylated DNA (120 sec), washed in immobilization buffer (30 sec), and then dipped into pre-equilibrated solutions of 2.5 nM Cra dimer plus varied concentrations of FruK. Immobilization buffer was similar to the PYK buffer used in the FruK enzyme assay. For DNA loading, immobilization buffer contained 80 nM DNA. Protein solutions were kept on ice until added to final reaction tube, which was then allowed to warm to room temperature (~15 min). (Biolayer interferometry signals are highly temperature dependent.) All biolayer interferometry steps were carried out with shaking at 2,200 rpm.

The signal changes associated with protein binding were monitored as a function of time until a plateau was reached. Experiments were repeated on separate days with different Cra and FruK preparations. The FruK could contribute up to ~ 1/15^th^ of the final reaction volume; because the FruK storage buffer differed significantly from the Cra storage buffer and contained 20% glycerol, we performed control experiments comprising Cra (i) pre-incubated with equivalent volumes of FruK storage buffer and (ii) with added glycerol; no effects were observed on DNA binding. Prior control experiments showed that FruK alone did not bind immobilized DNA (Supplementary Figure 11 in ^25^).

As extensively discussed in prior publications^25,76^, any equilibrium binding event should be assessed to determine whether it is in the “stoichiometric” binding regime, the “equilibrium” binding regime, or intermediate between the two. The feature that differentiates these regimes is the relationship of the fixed component of the binding reaction (here, immobilized Cra-DNA) to the K_d_ for the binding event. If (and only if) the concentration of the fixed component is at least 10-fold less than the K_d_ value, the binding event falls in the equilibrium regime, and K_d_ can be determined by fitting the binding curve with Equation 1, which is derived using several simplifying assumptions:

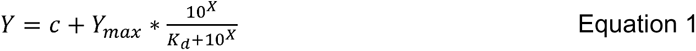

If the fixed component is *not* at least 10-fold below K_d_, then the binding reaction is in either the “stoichiometric” or “intermediate” regime and fits should use Equation 2 ^[83]^:

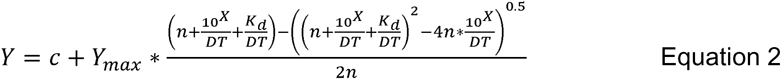

In both equations 1 and 2, “Y” is the observed signal, “c” is the baseline value when the concentration of protein is zero, “Y_max_” is the signal observed at saturation, “X” is the log of the FruK concentration (transformed to improve data point weighting in nonlinear regression), and “K_d_” is the equilibrium dissociation constant. In equation 2, “n” is the stoichiometry of the complex (here, n is assumed to 1) and “DT” is the total concentration of the species at fixed concentration.

The large number of parameters in equation 2 requires global fitting of multiple experiments to ascertain a reliable estimate of K_d_. In the BLI assays, we previously ascertained that the effective concentration of immobilized DNA (and therefore of DNA-bound Cra) is ~10^−9^ M^[25]^. Thus, for binding events with K_d_ values < 10^−8^ M, titration curves should be fit with Equation 2. Since the K_d_ value is not known at the start of an experiment, the binding regime must be determined from the shape of the binding curve and the effects of varying the fixed component (here, the concentration of immobilized DNA/Cra) in the binding reaction. For the Cra-FruK interaction, these processes revealed that the affinity was in the intermediate regime, with a K_d_ of 2.0 ± 0.5 x 10^−9^ M (Figure 4).

A second implementation of the BLI assay attempted to measure FruK binding to DNA-free Cra, using His-tagged Cra and FruK immobilized to Ni-NTA Dip and Read Biosensors (Forté Bio 18-5101; see Supplemental Results).

### Atomic force microscopy

For AFM, a support substrate of freshly cleaved mica was functionalized with a 167 μM solution of 1-(3-aminopropyl)-silatrane, as described previously^84^. Cra and FruK protein samples were mixed with DNA at the indicated concentrations in 1x Cra Buffer (50 mM HEPES, 100 mM KCl, 0.1 mM EDTA, 10 mM MgCl_2_, 0.3 mM DTT, pH 7.0) and incubated for 60 minutes at room temperature. After incubation, a 1:4 dilution of each sample was made using 1x Cra Buffer, of which 9 μL were immediately deposited on the functionalized mica surface. Deposited samples were incubated for 2 minutes, then rinsed with 200 μL ddH_2_O and dried with a gentle stream of argon. A typical image of 1×1 μm^2^ with 512 pixels/line was obtained under ambient conditions with a MultiMode AFM system (Bruker) using TESPA probes (Bruker Nano, Camarillo, CA, USA) housed at the Nanoimaging Core Facility at the University of Nebraska Medical Center. AFM images available to download from GitHub with the analyzed complexes indicated with boxes.

To analyze the particles observed in the AFM images, the volume of DNA-bound and DNA-free protein complexes were measured using AtomicJ^85^. The profile tool was used to obtain the following measurements: the complex height (h), the width at half-height of the long axis (w_long_), and the width at half-height of the orthogonal axis (w_orth_). For DNA-bound complexes, the height (h_DNA_) and width at half-height (w_DNA_) were also obtained for DNA adjacent to each DNA-bound complex. To calculate the volume (V) from the AFM images, peaks corresponding to protein complexes were fit as an elliptic cylinder

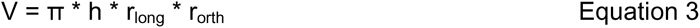

with the radii (r_long_ and r_orth_) being half of the measured widths. DNA was considered to be a cylindrical segment:

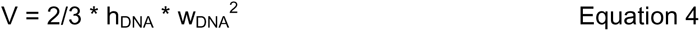

For DNA-bound complexes, the volume of adjacent DNA was subtracted so that only the volume attributed to the protein(s) remained. (Two other geometric models were considered for the protein peaks, but neither could be matched to the expected molecular weights: (i) Treating the peaks as a segment of a sphere under-estimated the volumes compared to expected molecular weights. (ii) Treating the peaks as a rectangular box over-estimated the volumes compared to expected molecular weights.)

To convert volumes to molecular weight, the nm^3^ volumes were divided by a conversion factor for the density. A value of 0.67 mL/g was determined from the crystal structures of both the Cra regulatory domain (PDB 2IKS;^32^) and ribokinase (PDB 1RKA^86^; ribokinase is a homolog of FruK). The density values were obtained by dividing the solvent-excluded volume (calculated by the *surface* command within UCSF Chimera^87^) by the molecular mass of the atoms present (mass for missing residues was subtracted). On the plots, reference lines for higher order complexes were calculated from the theoretical molecular weights for monomeric Cra (37,999 Da) and monomeric FruK (33,756 Da).

### Growth curves of WT and ΔfruK E. coli

To detect the biological outcomes of *fruK* deletion, we performed growth assays with the parent strain BW25113 (F-, Δ*(araD-araB)567,* Δ*lacZ4787*(::rrnB-3), λ*-, rph-1,* Δ*(rhaD-rhaB)568, hsdR514*) and its isogenic Δ*fruK724::kan* mutant JW2155-1 from the Keio collection^49^ that were obtained from the *E. coli* Genetic Stock Center at Yale University (New Haven, CT). To confirm the disruption of *fruK* in JW2155-1, the following primers were designed to flank the *fruK* gene (NCBI gene ID: BW25113_RS11315; WP_000091263.1): fwd 5’-agcagcagcgttttcattatg-3’; rev 5’-gacgctatcgctgctgg-3’. NEB Q5® High-Fidelity polymerase (Ipswich, MA) was used in a standard PCR reaction and the transposon disruption of *fruK* resulting amplicons was confirmed by sequencing (ACGT, Germantown, MD).

pHG165a (vector) and pHG165a-fruK (from EHEC) were transformed into BW25113 (wild type, WT) or Δ*fruK724*::*kan* (Δ*fruK*). Strains were grown overnight in “M9” minimal media (48 mM sodium phosphate, 22 mM potassium phosphate monobasic, 18.7 mM ammonium chloride, 8.5 mM sodium chloride, 2 mM magnesium sulfate, 0.1 mM CaCl_2_) supplemented with 0.2% glucose and ampicillin (100 ug mL^−1^) at 37°C with 250 rpm. The following day, cultures were pelleted at 4500xg using the Sorvall ST 16R Centrifuge for 10 minutes then washed twice with M9 minimal media with no carbon source added. The cells were then resuspended in M9 minimal media with no carbon source. The optical density was taken at a wavelength of 600 nm using Genesys 10S UV-Vis Spectrophotometer and adjusted to OD_600_ = 0.025 in 1mL of M9 media with either 0.2% glucose, fructose, or glycerol supplemented with ampicillin (100 μg mL^−1)^. These suspensions were then transferred into a nontreated 96 well plate in triplicate with 200 μL per well and incubated for 24 hours in a Tecan plate reader at 37°C with orbital shaking. OD600 readings were taken every 30 minutes.

A second version of the growth assay was modified from the protocol of Hall *et al.*^88^ and used uncomplemented cells with switching from LB to M9 media supplemented with 9 mM MgCl_2_. These Methods and Results are reported in Supplemental Figure 8. Results from the two implementations of the growth assay are in good agreement.

### Biofilm Assay

When assessing biofilm formation in different carbon source conditions, overnight cultures of the uncomplemented wild-type Keio and Δ*fruK724::kan* strains were grown in 3 mL LB media. Then cultures were spun down at room temperature for 5 min at 4,500xg (5,000 rpm) and the pellets were washed with 5 mL 1X MOPS (Teknova, M2101) followed by another spin at 4,500xg for 5 minutes. Pellets were resuspended with 2 mL 1X MOPS and split into 1.5 mL centrifuge tubes. These cultures were spun down at room temperature for 5 minutes at 10,621xg (10,000 rpm) and pellets were resuspended with 1mL, 1X MOPS with supplements (1X AUGC (Teknova, M2103), 1.23 mM KHPO_4_ (Teknova, M2102), 1X EZ solution (Teknova, M2104) and 0.2% glucose (Teknova, G-5802) or glycerol (Fisher-56-81-5) added buffer. Next, the resuspended culture was diluted to an OD_600_ of 0.1; 200 μL diluted culture was plated in a 96 sterile well plate (Fisher brand FB012931) and incubated at 30 °C for ~72 hours with saturating humidity.

At the end of the incubation, 75 μL of planktonic cell culture was transferred into a new plate and diluted with 75 μL fresh media. Planktonic growth was measured at OD_600_ to ensure that cultures reached comparable levels. The remaining media was removed from the growth plate, and wells were washed with autoclaved water to remove any remaining planktonic cells. 200 μL of crystal violet (Sigma-C6158; 0.1% in 12% ethanol) were added to the wells and incubated at room temperature for 15 minutes. After incubation, the dye solution was removed and wells were washed with ddH_2_O. Adherent dye was dissolved in 33% acetic acid and quantified by measuring the OD_595_.

## Acknowledgements

We thank the following KUMC scientists for their assistance with this work: Dr. Dipika Singh (initial FruK enzyme assays), Dr. Max S. Fairlamb (initial BLI assays), Mr. Benjamin F. Rau (assistance in the early stages of the project); and Dr. Alexey Ladokhin (English translation of ^47^). We thank the van Schaftingen laboratory for the gift of the *fruK* expression plasmid. Drs. Yuri Lyubchenko and Alexander Lushnikov at University of Nebraska Medical Center Nanoimaging Core Facility provided assistance with the AFM. This project was supported by the National Institute of General Medical Sciences at the National Institutes of Health by the following grants: the Center for Biomedical Research Excellence in Protein Structure and Function (P30 GM110761) to LSK, GM079423 to LSK, NIGMS P20 GM130448 to PRH, and GM113117 to PSH. This research was also supported by funding from the National Institute of Allergy and Infectious Diseases (NIAID) awards R01AI121073 and R21AI153773 to JLB and by private funds.

## Author contributions

LSK, PRH, JLB, SS, SEQ, and AWF contributed to the conception and design of the study; CW, CM, PTO, SEQ, KSH, SM, CLB, SS, and PSH contributed to the acquisition, analysis, and interpretation of the data; LSK, CW, JLB, AWF, PRH, SEQ, CM, and PTO contributed to writing the manuscript.

## Data Availability

The original AFM data files, a PDF identifying the analyzed particles, a copy of the AFM methodology, and an Excel file of the measurements collected are available online via GitHub: https://github.com/ptoneil/2022_LSK_AFM_CraFruK.git; all other data are in the main manuscript or the supplement.

## Supplemental Material

Supplemental Figures, Tables, Methods, Sequences, and Results can be found in the Supplemental Information. Additional citations associated with the supplemental material^89–93^.

