## Supplemental Information for "Fructose-1-kinase has pleiotropic roles in *Escherichia coli*"

**Supplemental Figure 1. *E. coli m*etabolite concentrations previously measured by Kochanowski et al.**^1^**.** (Left) Concentrations of both F-1-P and F-1,6-BP relative to their glucose M9 conditions were determined in >20 growth conditions. The concentrations of these two metabolites and were highly correlated except when fructose was the carbon source (right-most gray dot). The absolute concentrations of F-1,6-BP ranged from 0.03 to 3.7 mM; the absolute concentration of F-1-P could not be measured. (Right) The absolute concentrations of ATP and ADP under the same growth conditions.

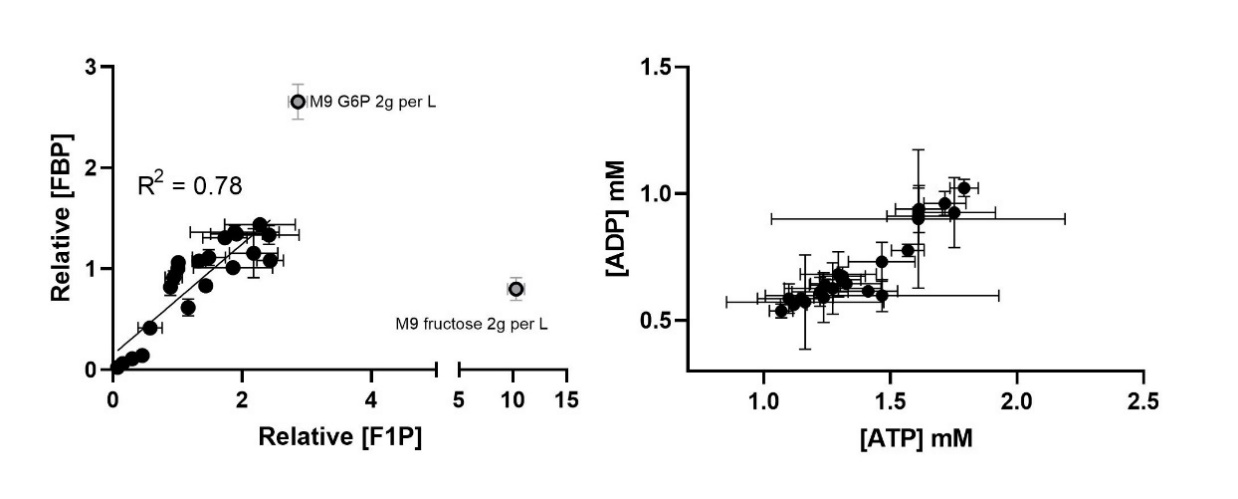

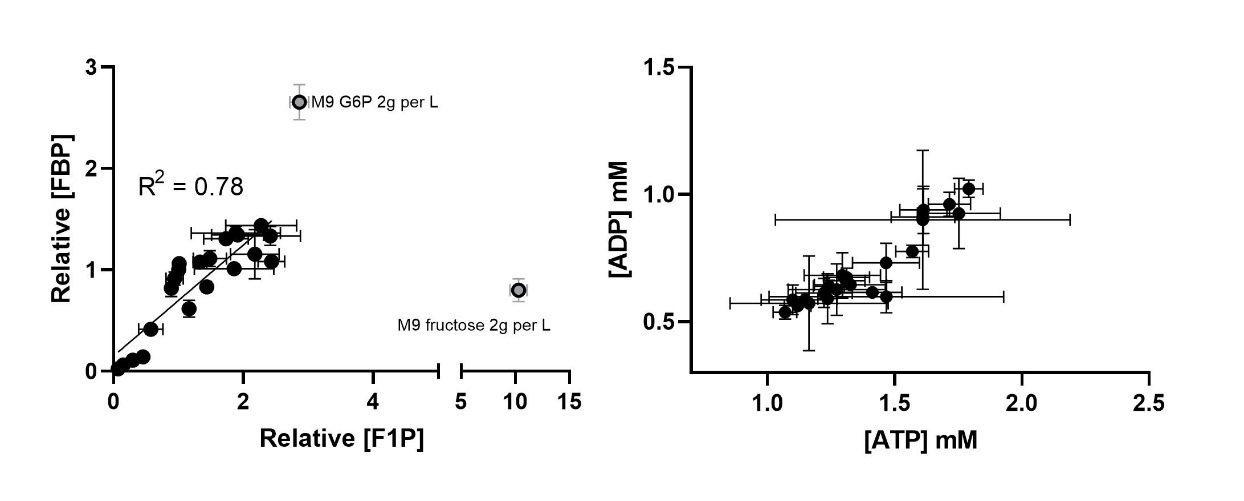

**Supplemental Table 1. Physiological ranges for FruK substrates in *E. coli* determined by Bennet *et al.* ^a^**

|  | Carbon source (4 g/l in minimal media) | | |
| --- | --- | --- | --- |
|  | glucose | glycerol | acetate |
| ATP | 9.6 mM^b^ | 9.0 mM^b^ | 4.1 mM |
| ADP^c^ | 0.55 mM | 0.15 mM | 0.19 mM |
| F-1,6-BP^d^ | 15 mM | 5.9 mM | < 0.1 mM |

^a^Values taken from Bennett *et al*.^2^. The concentration of F-1-P was not determined in this study.

^b^ATP differences between glucose and glycerol growth conditions were not statistically significant. ATP concentrations were also measured by Kochanoski *et al.* and ranged from 1 to 2 mM across >20 growth conditions (Supplemental Figure 1)^1^.

^c^ADP differences between the three growth conditions were not statistically significant. ADP concentrations were also measured by Kochanoski et al and ranged from 0.5 to 1 mM across >20 growth conditions (Supplemental Figure 1)^1^.

^d^The Kochanowski growth condition that matched most closely to that of Bennett *et al*. was 2g/L glucose^1^.

**Supplemental Figure 2. The “forward” FruK reaction converts F-1-P to F-1,6-BP.**

(Left) The forward FruK reaction was monitored by coupling ADP production to pyruvate kinase and lactate dehydrogenase and following the depletion of NADH at 340 nm^[3]^; components added to the reaction mixture are highlighted in yellow. (Right) Results from a representative assay; the units of the Y axis are ΔA_340nm_/min. The value reported for “Vmax” has *not* been normalized to FruK activity; the value reported for “Km” has units of mM.

The buffer and coupled reagents comprised: 20 mM HEPES pH=7.1, 5 mM MgCl_2_, 50 mM KCl, 0.5 mM DTT, 3 mM ATP, 0.33 mM NADH, 10 mM phosphoenolpyruvate (PEP), lactate dehydrogenase (Calzyme 092B0350, 275U/ml), and rabbit muscle pyruvate kinase (Roche 10128155001, 18U/ml). To balance the counter ions, F-1-P was diluted in 20 mM HEPES, pH 7.1, 5 mM MgCl_2_, 50 mM KCl. FruK was both stored and “washed” (see main text Methods) in buffer containing 20% glycerol; in the absence of glycerol, FruK activity quickly decays.

For both “unwashed” and “washed” FruK, K_m, F-1,6-BP_ values measured for multiple protein preparations have ranged from 0.9 to 3.5 mM.

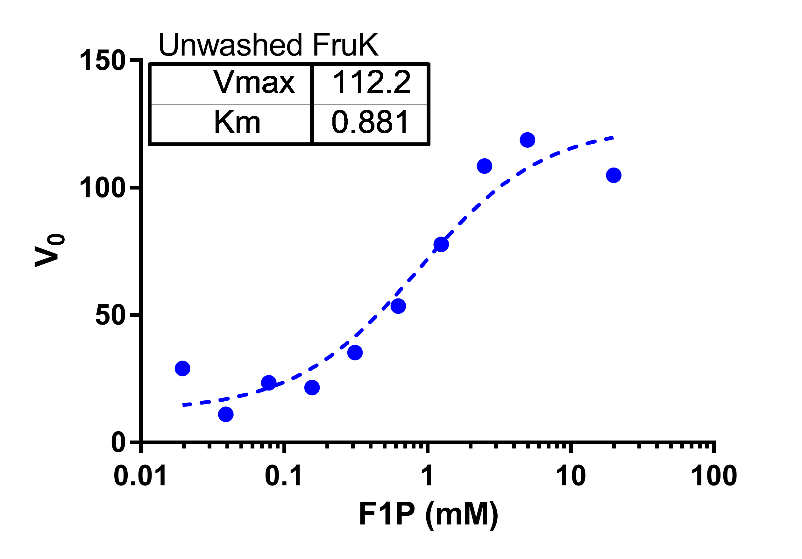

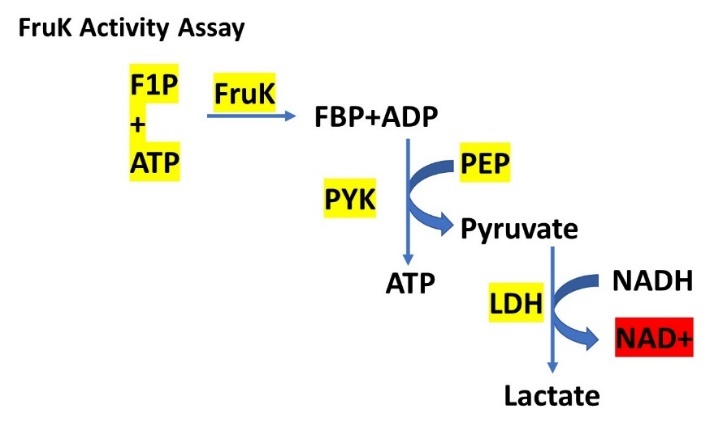

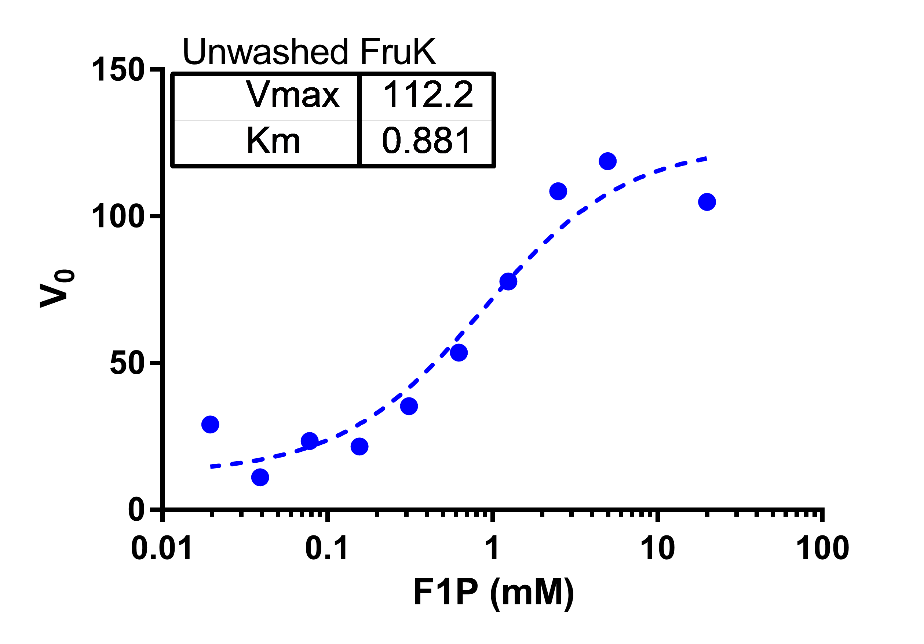

**Supplemental Figure 3. Domain Organization of Cra, LLhF and LacI.**

(Top) Schematic of functional domains in the LacI/GalR homolog monomer and dimer. Cra is a homolog in the LacI/GalR family. In these proteins, the DNA binding domain (oval) is joined to the regulatory domain (rounded rectangle) by an ~18 amino acid linker (bar)^4^. A repressor dimer is required to bind DNA; each monomer can bind one allosteric ligand (orange star) in a cleft on the regulatory domain.

(Bottom) Domain composition of *E. coli* LacI, chimeric LLhF*, and *E. coli* Cra. LLhF comprises the LacI DNA binding domain, the LacI linker, and the Cra regulatory domain. LLhF binds *lacO* DNA and responds to Cra’s allosteric ligands^4^. LacI dimers can dimerize to create a tetramer via a C-terminal Leucine-heptad repeat sequence (horizontal bars in the bottom left cartoon), which allows looping if two DNA operators are on the same piece of DNA^5^; this C-terminal domain is not present in Cra or LLhF.

*The name “LLhF” derives from the fact that original name for Cra was “FruR” (abbreviation for “fructose repressor protein”); the name was changed when the extent of the Cra regulon became known^6^.

**Supplemental Results: Wild-type Cra pulls-down FruK only when cells are grown in fructose.**

**Allosteric ligand**

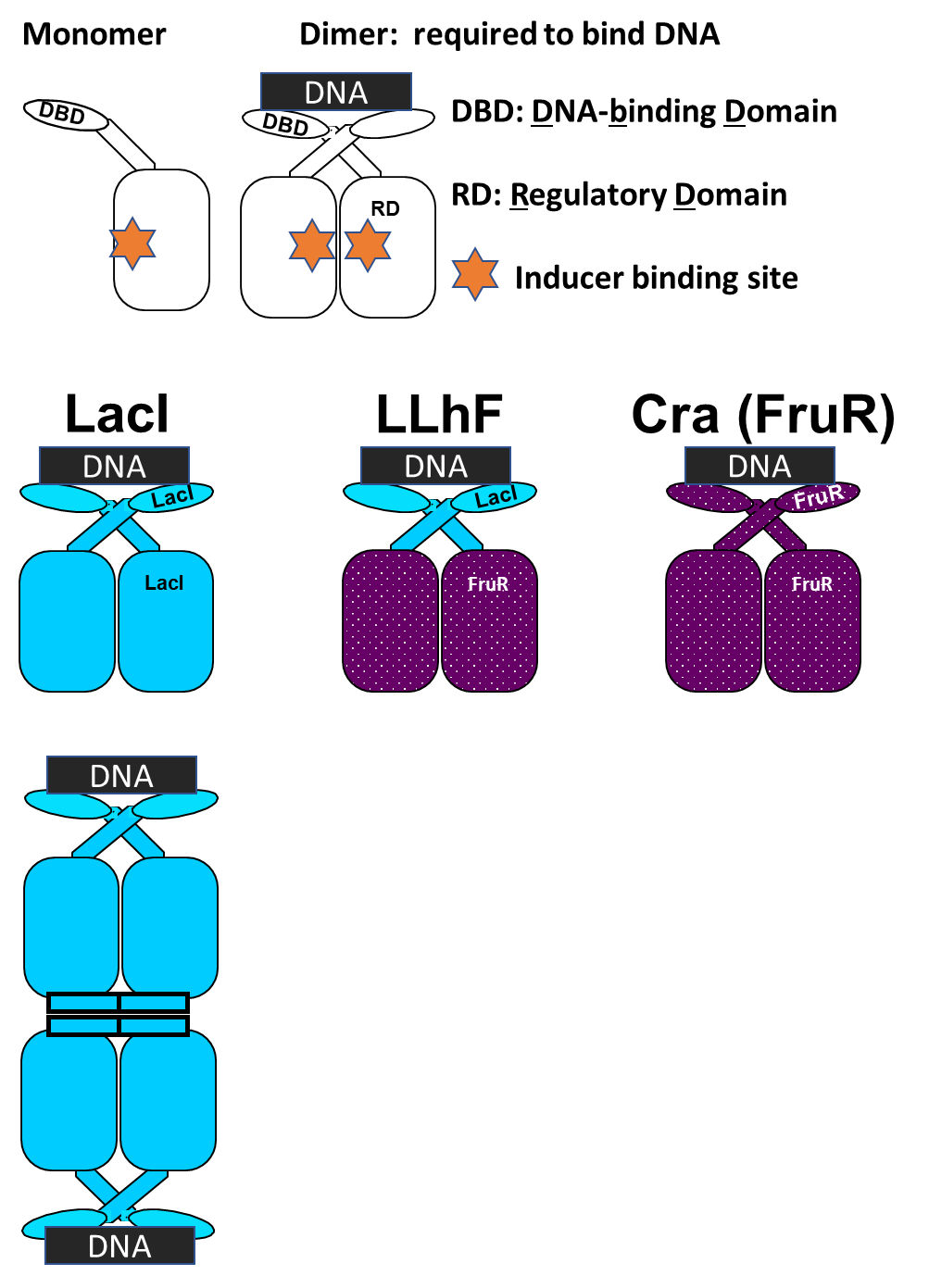

**RD**

The pull-down data shown in Figure 3 support the hypothesis that FruK interacts with full-length Cra, but the fact that the interaction is only observed when cells were growth in 20 mM fructose raises two confounding observations that must be resolved.

The first apparent conundrum is that *E. coli* should convert the 20 mM fructose to the strong Cra allosteric inhibitor F-1-P. In turn, this should *diminish* the Cra-DNA interaction required for the pull-down assay. However, prior to the pulldown, much of the F-1-P would be further metabolized during over-night growth of the culture and any remaining F-1-P should be washed away during sample preparation. Furthermore, even if substantial amounts of F-1-P are present in the current experiment, it is reasonable that Cra maintains an ability to bind DNA in the pull-down assay: Cra is a homolog of LacI; although binding to its allosteric ligand diminishes LacI-DNA binding several orders of magnitude, “diminish” does not mean “abolish”. Various assays have to detected this weakened binding (*e.g.* ^3^) and LacI was visible in pull-down assays even in the presence of its allosteric inhibitor (data not shown). Likewise, >100 repressor variants with weak DNA binding (*i.e.* not tight enough to facilitate repression) were successfully visualized using this DNA pull-down assay^7^.

A second apparent conundrum of the pull-down results is to explain why engineered LLhF and wild-type Cra required different growth conditions to show the FruK interaction. An important clue is that Cra represses the *fruBKA* operon. As such, very little FruK would be expressed in the MOPS-glycerol media, making it difficult to detect in SDS-PAGE. However, expression levels would rise when the *fruBKA* operon was de-repressed. For cells constitutively over-expressing Cra, de-repression was accomplished by the conversion of fructose to F-1-P (before its further metabolism). In cells constitutively over-expressing the engineered LLhF, *fruBKA* de-repression could be accomplished *via* formation of dominant negative heterodimers with endogenous wild-type Cra. As precedence for this hypothesis, overexpression of LacI:PurR and LacI:GalR chimeras de-repress genes in their regulons (unpublished data). These three chimeras use, respectively, the Cra, PurR, and GalR regulatory domains, which almost exclusively mediates dimer formation in the LacI/GalR paralogs^5^ and should be able to form hetero-dimers with their wild-type, endogenous counterparts.

**Supplemental Figure 4. Specificity of the Cra-DNA FruK interaction**. Biotinylated operator DNA was immobilized to strept­a­vidin sen­sor tips for BLI assays. (Left) 100 nM Cra was bound to immobilized operator “FruB-prox”; the addition of 3 μM FruK increased signal, indicative of binding. As previously shown, when the Cra-FruK interaction reaches equilibrium, the height of the plateau correlates with FruK concentration and can be used to construct the titration plots shown in Figure 4 of the main text. In control experiments, purified *E. coli* pyruvate kinase (“PYK”; 1.3 µM; ^8^) does not bind DNA-Cra and acts as a specificity control; other PYK concentrations showed no binding. Note that neither FruK nor PYK binds to immobilized DNA in the absence of Cra.

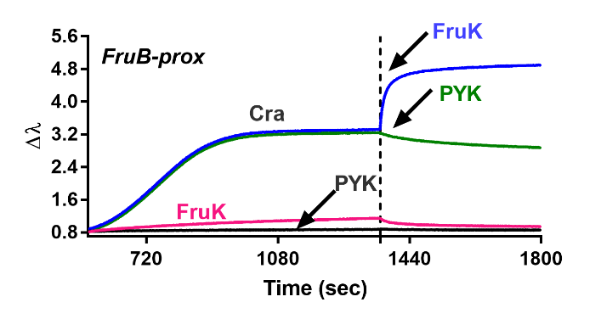

**Supplemental Figure 5**. **Atomic force microscopy.**

All raw AFM data files for each sample, as well as images showing selected particles, are available for download at <https://github.com/ptoneil/2022_LSK_AFM_CraFruK.git>. Each image is displayed using the same depth scale from 0.2 to 1 nm; the actual maximum depth for most AFM images exceeded 3 nm. Each of the widefield views shows a 1x1 µm^2^ AFM image unless otherwise indicated. Boxes (50x50 nm^2^ unless otherwise indicated) indicate the complexes that were analyzed. Cra and FruK concentrations are provided as M dimer.

The image below shows that FruK does not directly bind DNA. A sample comprising 20 nM FruK and10 nM DNA was diluted 1:4 prior to adherence on functionalized mica (2x2 µm^2^ AFM image). Most DNA appears as in the DNA-only sample, with no significant FruK binding. Instead, multiple densities were observed in the background, which likely corresponds to free FruK. This behavior is consistent with other observations that FruK lacks the ability to bind to DNA. Each box (100x100 nm^2^) indicates an analyzed, DNA-free protein complex.

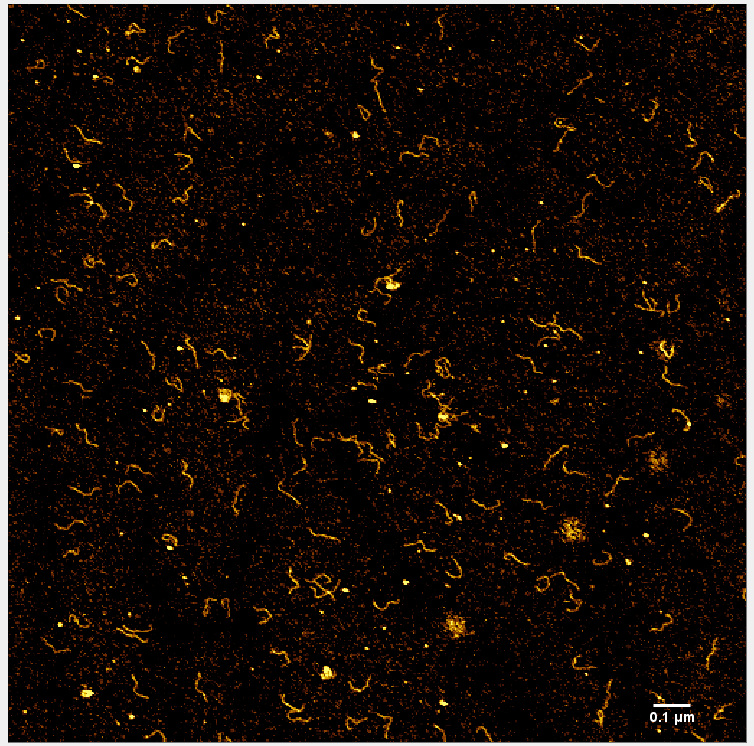

**Supplemental Figure 6. The Cra/FruK ratio affects the population of higher order complexes in AFM.** A. Comparison DNA-bound and DNA-free Cra + FruK populations at two different Cra/FruK ratios. For samples containing the 20 nM Cra and 10 nM FruK, the population showed a higher fraction of large complexes than did the sample containing 40 nM each Cra and FruK. This suggests that the higher FruK concentration inhibited formation of higher order complexes, perhaps by interrupting the C_4_F_2_ complex with additional F_2_ (see B, below)

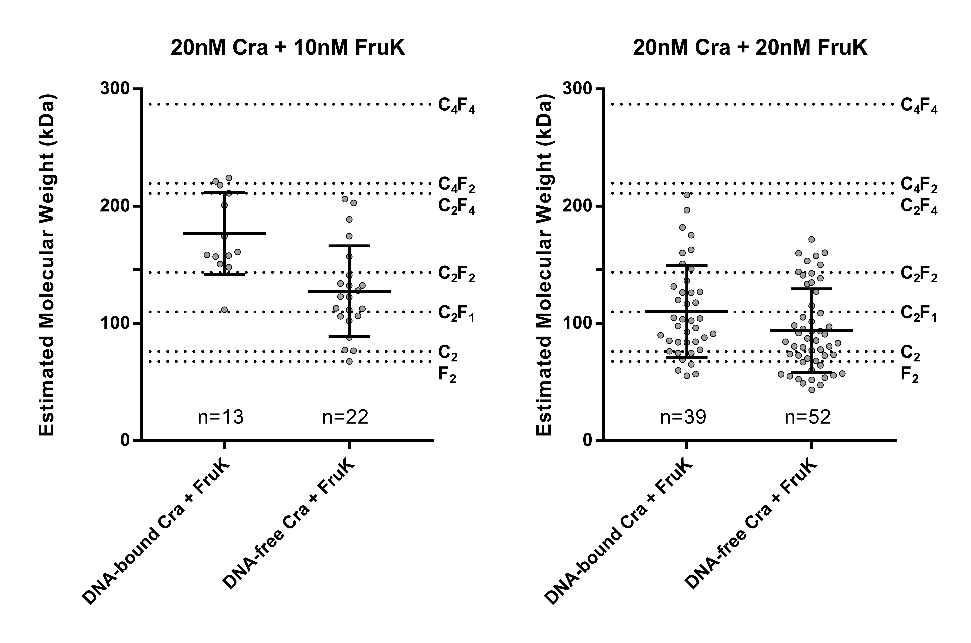

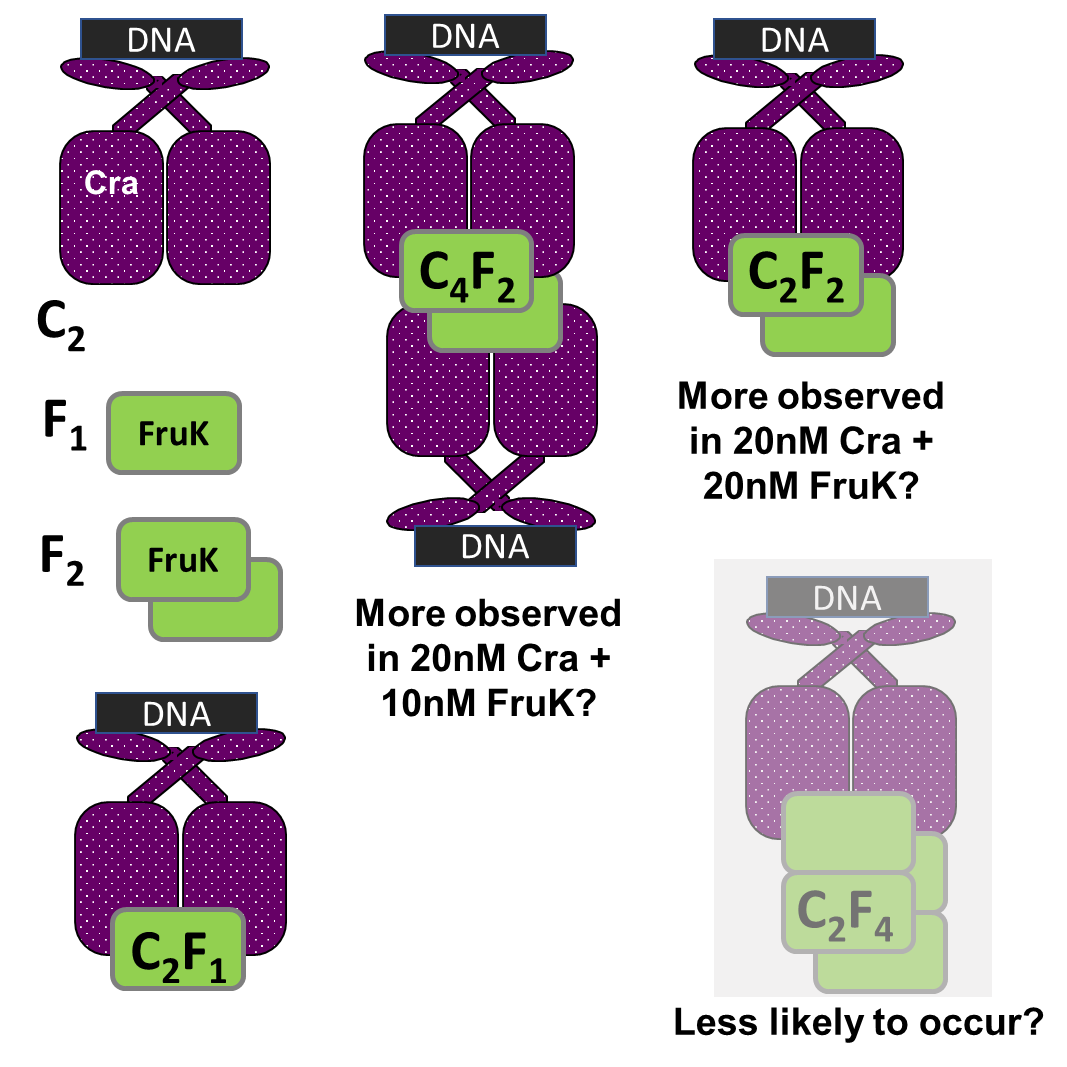

(dimer)_1_

(dimer)_2_

(dimer)_3_

B. Cartoon schematics of Cra/FruK complexes, along with notations as to which may correspond to (dimer)_1_, (dimer)_2_, and (dimer)_3_ in the AFM.

(dimer)_1_

**Supplemental Figure 7**. **Alternative clustering of the DNA-bound Cra plus FruK population.** On the plots below, each dot represents a unique, measured peak in the AFM image; the bar represents the mean and standard deviation for each population. One-dimensional cluster analyses of the DNA-bound Cra + FruK population from Figure 5B were performed with two-, three-, or four-classes using “Jenk’s Natural Breaks”. This method evaluates the outcomes from separating all possible combinations of particles into the specified number of classes. Within a designated fit, the best-fit clustering is identified as having the minimum sum of subclass variance; to discriminate among fits, we compared the average MWs of each class to expected MWs of various Cra/FruK complexes (dotted reference lines; Cra: 37,999 Da; FruK: 33,756 Da).

In contrast to the three-class clustering shown in Figure 5C of the main text, several of the means determined by two- and four-class clustering (left and right-most panels) did *not* correlate with the expected molecular weights of higher order complexes.

For the unfixed, three-class analysis (middle panel), particles in two of the identified classes had mean MWs near the expected molecular weights for unbound Cra dimers (C_2_) and Cra dimer/FruK dimer (C_2_F_2_). However, the mean for the third (highest) molecular weight class was lower than expected, and the three lowest data points of this set subjectively appear misclassified. Misclassification could be due to the limited number of higher molecular weight complexes; when this number was artificially increased by duplicating 3 upper data points, the lower three data points were re-classified to class 2 (not shown).

Thus, for the three-class clustering shown in Figure 5C of the main text, the upper class was fixed at n=7, corresponding to the upper class in the unfixed, four-class clustering (right-most panel). As a control calculation, the same fixed cluster analysis was performed for the four-class system; no difference was observed.

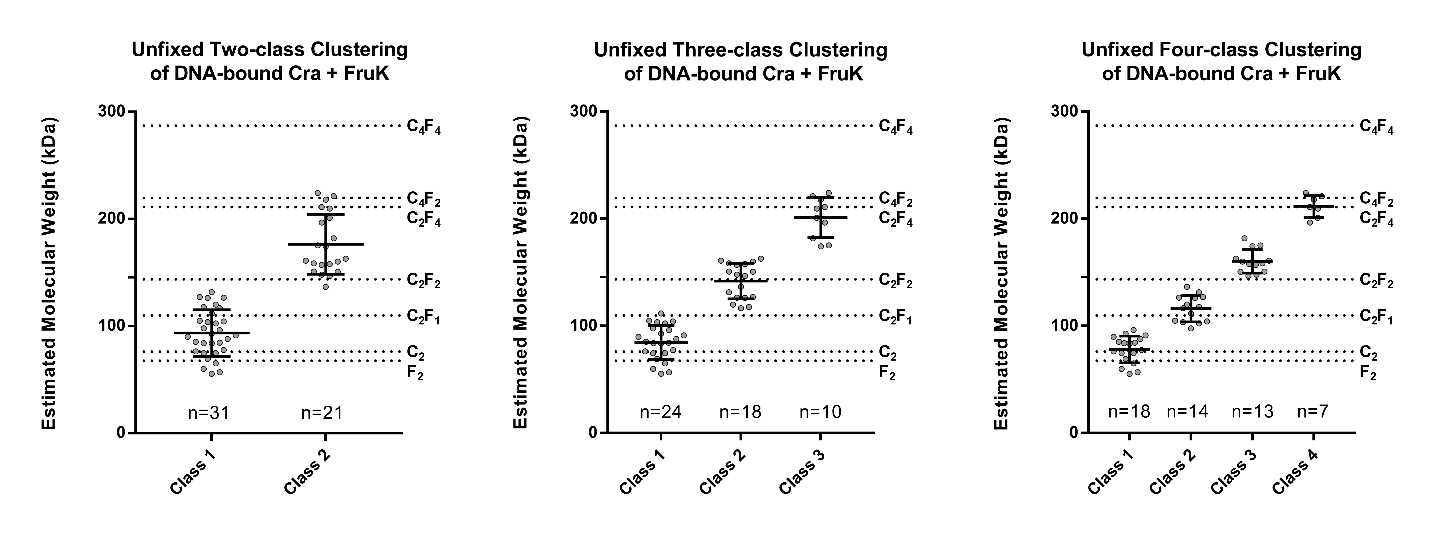

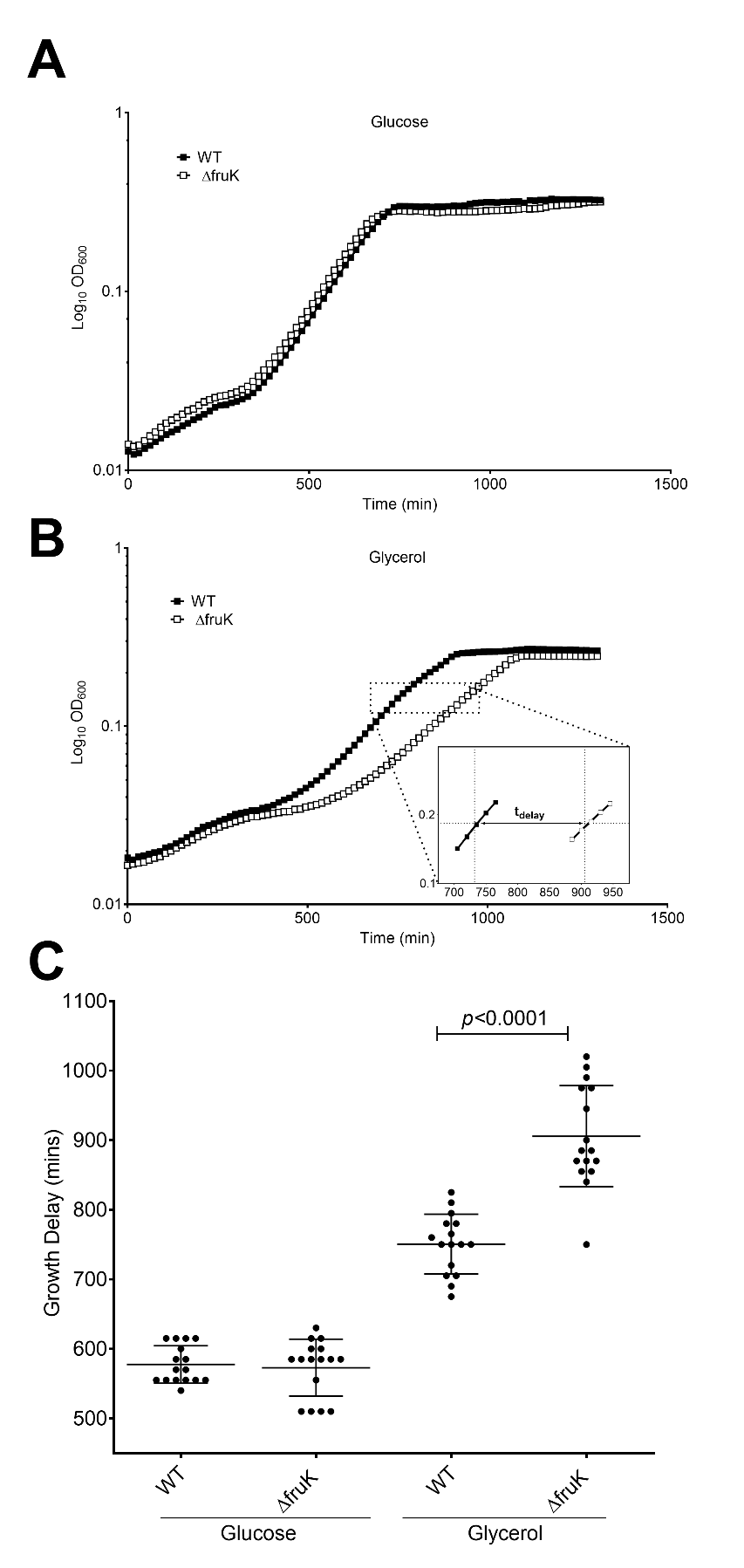
**Supplemental Figure 8.** **Growth of WT and *ΔfruK::kan* (“*ΔfruK*”) strains when shifted from LB to M9 defined media, (no comple­men­ta­tion).** Wild-type and Δ*fruK::kan* strains were grown in 10 mL rich media (Luria broth) until late-log phase (OD600~1.5 in a cuvette with 1 cm path length). Cultures were then centrifuged at 1800xg for 15 mins, decanted, and resuspended in 1mL M9 media (42 mM sodium phosphate, 22 mM potassium phosphate monobasic, 18.7 mM ammonium chloride, 8.6 mM sodium chloride, 2 mM magnesium sulfate, 0.02 mM CaCl_2_, 0.00005% thiamine, pH 7.4) containing 9 mM MgCl_2_ and 0.2% w/v of glucose, glycerol, or fructose as the carbon source, to a starting OD_600_ of ~0.1. Subsequently, 100µL of resuspended bacteria were aliquoted into quadruplicate wells of a 96-well plate (Becton Dickinson, NJ).

Bacterial growth was then monitored using either a BioTek Synergy2 microplate reader (Winooski, VT) or a Tecan M200 pro (Switzerland) with data collection *via* either Gen5 or Magellan data analysis software. The plate was kept at 37°C with shaking (198.4 rpm); OD_600_ was measured every 15 mins for 24 hours. Media-only wells served as blanks and were subtracted from bacteria-containing wells. Doubling times were calculated for the logarithmic growth phase using the inverse slope of log_10_(OD_600_) values. Data shown are the mean and standard deviation of quadruplicates samples from each of four independent experiments; unpaired t-tests using were used to assess differences in growth rates. The growth delay time as the wild-type and Δ*fruK::kan* strains transitioned from rich to minimal media (t_delay_) was calculated as the mean time required for bacteria to reach mid-log phase (OD_600_ 0.15).

Growth in (A) glucose and (B) glycerol. Delay time (t_delay,_ insert B) was calculated as the amount of time to reach mid-log phase and in (C) is shown as mean and standard deviation of quadruplicate wells from four separate experiments. In glycerol, the mutant showed a significantly increased delay as compared to WT (*p*<0.0001, paired t-test).

**Supplemental Methods:** **Preparation of DNA for AFM.**

For AFM, a 331 bp of region of the *fruBKA* promoter, spanning both the *fruB_distal_* and *fruB_proximal_* operator binding sites, was amplified from the DH5α strain of *E. coli*. To prepare genomic DNA, log-phase *E. coli* grown in LB were centrifuged, resuspended in 1/5 volume milliQ water, and boiled at 95°C for 10min. Amplification of the promoter region was carried out using primers with added restriction sites (underlined):

fruBKA_cra_EcoRI_For: 5’-atgaattcgcgaatcgcctcttctttgtctc and

fruBKA_cra_HindIII_Rev: 5’-ataagcttgctgacgctgctgaaatgaaattgc;

the restriction sites allow for future subcloning into a plasmid.

Amplification reactions contained 5ul 10x PfuUltraBuffer (Agilent), 1.25uL 10mM dNTPs, 2ul (~100ng) genomic DNA, 1uL 10uM each primer, and 1uL PfuTurbo (Agilent). A two-phase amplification comprised annealing at 53.2°C for 10 cycles and then at 56.4°C for 25 cycles. The amplification mixture was analyzed with agarose gel electrophoresis and showed a single band with the expected molecular weight. Fragment sequencing (ACGT) with the amplification primers confirmed the expected sequence (below).

In the sequence below, the green highlight indicating the start codon for *fruB*. The bold regions indicate the *fruB_distal_* and *fruB_proximal_* operator sites (of the pair, *fruB_proximal_* is highlighted in purple); underlined regions indicate the engineered restriction enzyme sites.

5’‑ATAAGCTTGCTGACGCTGCTGAAATGAAATTGCTGATAACGGATTTTCCCATCAGCAATTAGGAAAAATGGCAAAAAATTGTGCAGCACATCAAACTTTTGCTCATAACTTTACGGCTTTCCTTGCGTGCTGAAACAGAAGTATTATGC**TTTCTTGAAACGTTTCAGC**GCGATCTTGTCTTTAACCCTAAGCCAGGTTGGCGCTTTTTTTCTCAT**AGAGGCTGAATCGTTTCAA**TTCAGCAAGAGAGGAGAACTATGTTCCAGTTATCCGTACAGGACATCCATCCGGGCGAAAAGGCCGGAGACAAAGAAGAGGCGATTCGCGAATTCAT

**Supplemental Table 2. Bacterial strains and plasmids used in this study.**

| **Strain name** | **Relevant characteristics** | **Use** |
| --- | --- | --- |
| 3.300 ^9^ | Hfr(PO1)*, lacI22, λ-, e14-, relA1, spoT1, thiE1* | Source of *cra* gene; DNA pull-down assays |
| BLIM ^10^ | *omp*T, *hsdS_B_*(*r_B_*^–^*m_B_*^–^)^-^, *gal, dcm, lac* | Expression of Cra protein |
| BL21 (DE3) pLysS | F–, *omp*T, *hsd*S_B_ (r_B_-, m_B_-), *dcm*, *gal*, λ(DE3), [malB^+^]_K-12_(λ^S^), pLysS, Cm^r^ | Expression of FruK protein |
| BL21 (DE3) | F –, *omp*T, *hsd*S_B_ (r_B_- m_B_-) *dcm, gal,* λ (DE3), [malB^+^]_K-12_(λ^S^) | FruK-Cra pull-down assay |
| DH5α | F^–^ φ80*lac*ZΔM15 Δ(*lac*ZYA-*arg*F)U169 *rec*A1 *end*A1 *hsd*R17(r_K_^–^, m_K_^+^) *pho*A *sup*E44 λ^–^*thi*-1 *gyr*A96 *rel*A1 | Source of *fruBKA* promoter for AFM |
| BW25113 ^11^ | Wildtype *E. coli* strain F-, *Δ(araD-araB)567, ΔlacZ4787*(::rrnB-3)*, λ-, rph-1, Δ(rhaD-rhaB)568, hsdR514* | Growth and biofilm assays |
| JW2155-1 ^11^ | BW25113 Δ*fruK724::kan* | Growth and biofilm assays |
| EDL933 ^12^ | O157:H7 enterohemorrhagic *E. coli* strain | Source of EHEC *cra* and *fruK* gene |
| **Plasmid name** |  |  |
| pHG165a ^4,13^ | Parent plasmid (vector) | Growth assays |
| FruR-pHG165a | Wild-type Cra; AddGene 90066; constitutive expression | Expression of untagged Cra protein |
| pET28a-Cra-C-Flag | Inducible expression | Expression of Cra-Flag protein |
| pET3a-Cra-C-His | Inducible expression | Expression of Cra-His protein |
| pET-3a-FruK ^14^ | Wild-type FruK; inducible expression | Expression of un­tagged FruK protein |
| pET28a-FruK-His | Inducible expression | Expression of FruK-His protein |
| pET3a-FruK-C-His | Inducible expression | BLI trials |
| pHG165a-FruK | Wild-type FruK complementation plasmid; constitutive expression | Growth assays |

**Supplemental Table 3.** Primers used to generate Cra and FruK expression plasmids

| **Primer** | **Sequence 5’-3’** |
| --- | --- |
| Primers used to clone *E. coli fruK* open reading frame into pHG165a and pET28a. | |
| pHG165a/ pET28a primer A1 | GTTACCTTCGGAAAAAGAGTTGGTAGCTCTTGATCCGG |
| pET28a primer A2 | GTATATCTCCTTCTTAAAGTTAAACAAAATTATTTCTAGAGG |
| pHG165a primerA2 | ATTCACCACCCTGAATTGACTCTCTTCCGGG |
| pET28a primer C1 | GGTGGCTCGGATGGTTCAACTAGTGGTTCTGGTCATCACCATCACCATCACTGAGATCCG |
| pHG165a primerC1 | TGAGAATTCGTAATAATATAGCTCTAGATTAATTGCG |
| pHG165a C ter His primer C1 | CAACTAGTGGTTCTGGTCATCACCATCACCATCACTGATGAGAATTCGTAATAATATAGC |
| pHG165a/ pET28a primer C2 | CCGGATCAAGAGCTACCAACTCTTTTTCCGAAGGTAAC |
| *fruK* in pHG165a primerB1 | CCGGAAGAGAGTCAATTCAGGGTGGTGAATATGAGCAGACGTGTTGCCACTATCACCCT |
| Untagged *fruK* primer B2 | CAATTAATCTAGAGCTATATTATTACGAATTCTCATCAGTTAAACGGTTGTAAGTCGACG |
| *fruK* C -ter His primer B2 | AACCATCCGAGCCACCTTTGCCACCATGGTCGCCACCACCGCCGTTAAACGGTTGTAAGT |
| *fruK* in pET28a primerB1 | GAAATAATTTTGTTTAACTTTAAGAAGGAGATATACCATGAGCAGACGTGTTGCCACTAT |
| *fruK* C -ter His in pET28a primer B2 | GTTGAACCATCCGAgccaccTTTgccaccATGGTCGCCACCACCgccGTTAAACGGTTGTAAG |
| Primers used to clone *E. coli cra* open reading frame into pHG165a. | |
| *cra* For | GGTCTCCAGGTGTGAAACTGGATGAAATCGC |
| *cra* Rev | GAGCTCTTAGCTACGGCTGAGCACGCCGCGG |
| Primers used to create His- and Flag-tagged versions of the *cra* and *fruk* coding regions. | |
| cra 6His For | CGCCGCGGCGTGCTCAGCCGTAGCCATCATCATCATCATCACTAATAACCTGAGGCCG |
| cra 6His Rev | CGACAGTATCGGCCTCAGGTTATTAGTGATGATGATGATGATGGCTACGGCTGAGCACG |
| cra c-ter flag for | CCGCGGCGTGCTCAGCCGTAGCCTACAAGGACGATGACGACAAGTGTAACCTGAGGCCG |
| cra c-ter flag rev | GACAGTATCGGCCTCAGGTTACACTTGTCGTCATCGTCCTTGTAGGCTACGGCTGAGCAC |
| 6His fruK for | GTTTAACTTTAAGAAGGAGATATACAtATGCATCATCATCATCATCACATGAGCAGACGT |
| 6His fruK rev | GGGTGATAGTAGCAACACGTCTGCTCATGTGATGATGATGATGATGCATaTGTATATCTC |
| fruK 6His For | GTCGACTTACAACCTTTTAACCATCATCATCATCATCACTGActgGATCCGGCTGCTAAC |
| fruK 6His Rev | CGGGCTTTGTTAGCAGCCGGATCcagTCAGTGATGATGATGATGATGGTTAAAAGGTTG |

**Supplemental Sequences**

A. The Cra sequence used in this work corresponds to Uni-Prot P0ACP1.

MKLDEIARLAGVSRTTASYVINGKAKQYRVSDKTVEKVMAVVREHNYHPNAVAAGLRAGRTRSIGLVIPDLENTSYTRIANYLERQARQRGYQLLIACSEDQPDNEMRCIEHLLQRQVDAIIVSTSLPPEHPFYQRWANDPFPIVALDRALDREHFTSVVGADQDDAEMLAEELRKFPAETVLYLGALPELSVSFLREQGFRTAWKDDPREVHFLYANSYEREAAAQLFEKWLETHPMPQALFTTSFALLQGVMDVTLRRDGKLPSDLAIATFGDNELLDFLQCPVLAVAQRHRDVAERVLEIVLASLDEPRKPKPGLTRIKRNLYRRGVLSRS

B. The FruK sequence used in this work, as translated from the *fruK* coding region of the pET3a plasmid from the Van Schaftingen lab (AddGene 186256) used for experiments represented in Figure 2, 3A, 4, and 5.

MSRRVATITLNPAYDLVGFCPEIERGEVNLVKTTGLHAAGKGINVAKVLKDLGIDVTVGGFLGKDNQDGFQQLFSELGIANRFQVVQGRTRINVKLTEKDGEVTDFNFSGFEVTPADWERFVTDSLSWLGQFDMVCVSGSLPSGVSPEAFTDWMTRLRSQCPCIIFDSSREALVAGLKAAPWLVKPNRRELEIWAGRKLPEMKDVIEAAHALREQGIAHVVISLGAEGALWVNASGEWIAKPPSVDVVSTVGAGDSMVGGLIYGLLMRESSEHTLRLATAVAALAVSQSNVGITDRPQLAAMMARVDLQPFN*

C. The FruK sequence from EHEC (strain EDL933), subcloned onto pHG165a/c and used for the protein-protein pull-down in Figure 3BC and Figure 6.

MSRRVATITLNPAYDLVGFCPEIERGEVNLVKTTGLHAAGKGINVAKVLKDLGIDVTVGGFLGKDNQDGFQQLFSELGIANRFQVVQGRTRINVKLTEKDGEVTDFNFSGFDVTPADWERFVNDSLSWLGQFDMVCVSGSLPAGVSPEAFTDWMTRLRSQCPCIIFDSSREALVAGLKAAPWLVKPNRRELEIWAGRKLPEMKDVIDAAHALREQGIAHVVISLGAEGALWVNASGEWIAKPPAVDVVSTVGAGDSMVGGLIYGLLMRESSEHTLRLATAVAALAVSQSNVGITDRPQLAAMMARVDLQPFN

D. Alignment of *E. coli* ribokinase and representative FruK sequences found in the nr database, including the sequence encoded on pET3a (Uniprot P0AEW9) and from EHEC, both used in this work. Yellow shading highlights differences among the FruK sequences. Differences between between Uniprot P0AEW9 (K12) and EHEC FruK sequences are bold and underlined. The differences in these two sequences have similar side chain properties and/or were observed in the FruK sequences from other *E. coli* strains.

```
Ribokinase(P0A9J6)	MQNAGSLVVLGSINADHILNLQSFPT---PGETVTGNHYQVAFGGKGANQAVAAGRSGAN	57
EHEC EDL933	-MSR----RVATITLNPAYDLVGFCPEIERGEVNLVKTTGLHAAGKGINVAKVLKDLGID	55
Uniprot_P0AEW9	-MSR----RVATITLNPAYDLVGFCPEIERGEVNLVKTTGLHAAGKGINVAKVLKDLGID	55
WP_025798139.1	-MSR----RVATITLNPAYDLVGYCAEIERGEVNRVQTAGLHAAGKGINVAKVLKDLGID	55
RDT52444.1	-MSR----RVATITLNPAYDLVGFTPEIERGEVNLVRTTGLHAAGKGINVAKVLKDLGID	55
WP_001653212.1	-MSR----RVATITLNPAYDLVGFCPEIERGEVNLVKTTGLHAAGKGINVAKVLKDLGID	55
EEZ4480454.1	-MSR----RVATITLNPAYDLVGFCPEIERGEVNLVKTTGLHAAGKGINVAKVLKDLGID	55
WP_163341112.1	-MSR----RVATITLNPAYDLVGFCPEIERGEVNLVKTTGLHAAGKGINVAKVLKDLGID	55

Ribokinase(P0A9J6)	IAFIACTGDDSIGESVRQQLATDNIDIT-PVSVIKGESTGVALIFVNGEGENVIGIHAGA	116
EHEC EDL933	VTVGGFLGKDNQD-GF-QQLFS-ELGIANRFQVVQGR-TRINVKLTEKDGEVTDFNFSGF	111
Uniprot_P0AEW9	VTVGGFLGKDNQD-GF-QQLFS-ELGIANRFQVVQGR-TRINVKLTEKDGEVTDFNFSGF	111
WP_025798139.1	VTVGGFLGKENQD-GF-QQLFS-ELGIANRFQVVAGR-TRINVKLTEKDGEVSDFNFSGF	111
RDT52444.1	VTVGGFLGKDNQD-GF-QQLFS-ELGIANRFQIVQGR-TRINVKLTEKDGEVTDFNFSGF	111
WP_001653212.1	VTVGGFLGKDNQD-GF-QQLFS-ELGIANRFQVVQGR-TRINVKLTEKDGEVTDFNFSGF	111
EEZ4480454.1	VTVGGFLGKDNQD-GF-QQLFS-ELGIANRFQVVQGR-TRINVKLTEKDGEVTDLNFSGF	111
WP_163341112.1	VTVGGFLGKNNQD-GF-QQLFS-ELGIANRFQVVQGR-TRINVKLTEKDGEVTDFNFSGF	111

Ribokinase(P0A9J6)	NAALSPALVEAQRERIANASA-LLMQLE-----SPLESVMAA-------AKIAHQNKTIV	163
EHEC EDL933	------DVTPADWERFVNDSLSWLGQFDMVCVSGSLPAGVSPEAFTDWMTRLRSQCPCI-	164
Uniprot_P0AEW9	------EVTPADWERFVTDSLSWLGQFDMVCVSGSLPSGVSPEAFTDWMTRLRSQCPCI-	164
WP_025798139.1	------EVTKQDWERFVNDSLSWLGQFDMVCVSGSLPAGVDPDDFTDWMRRLRSQCPCI-	164
RDT52444.1	------EVTPADWERFVNDSLTWLGQFDMVCVSGSLPSGVSPEAFTDWMTRLRSQCPCI-	164
WP_001653212.1	------EVPPADWERFVTDSLSWLGQFDMVCVSGSLPSGVSPEAFTDWMTRLRSQCPCI-	164
EEZ4480454.1	------EVTPADWERFVTDSLSWLGQFDMVCVSGSLPSGVSPEAFTDWMTRLRSQCPCI-	164
WP_163341112.1	------EVTPADWERFVTDSLSWLGQFDMVCVSGSLPSGVSPEAFTDWMTRLRSQCPCI-	164

Ribokinase(P0A9J6)	ALNPAPARELPDELLALVDIITPNETEAEKLTGIRVENDEDAAKAAQVLHEKGIRTVLIT	223
EHEC EDL933	-IFDSSREALVAGLKAAPWLVKPNRRELEIWAGRKLPEMKDVIDAAHALREQGIAHVVIS	223
Uniprot_P0AEW9	-IFDSSREALVAGLKAAPWLVKPNRRELEIWAGRKLPEMKDVIEAAHALREQGIAHVVIS	223
WP_025798139.1	-IFDSSREALVAGLKAAPWLVKPNRRELEIWAGRELPTLDDVVGAAHALRDQGIAHVVIS	223
RDT52444.1	-IFDSSRDALVAGLKAAPWLVKPNRRELEIWAGRKLPELKDVIDAAHALREQGIAHVVIS	223
WP_001653212.1	-IFDSSREALVAGLKAAPWLGKPNRRELEIWAGRKLPEMKDVIEAAHALREQGIAHVVIS	223
EEZ4480454.1	-IFDSSREALVAGLKAAPWLVKPNRRELEIWAGRKLPEMKDVIEAAHALREQGIAHVVIS	223
WP_163341112.1	-IFDSSREALVAGLKAAPWLVKPNRRELEIWAGRKLPEMKDVIEAAHALREQGIAHVVIS	223

Ribokinase(P0A9J6)	LGSRG-VWASVNGEGQRVPGFRVQAVDTIAAGDTFNGALITALLEEKPLPEAIRFAHAAA	282
EHEC EDL933	LGAEGALWVNASGEWIAKP-PAVDVVSTVGAGDSMVGGLIYGLLMRESSEHTLRLATAVA	282
Uniprot_P0AEW9	LGAEGALWVNASGEWIAKP-PSVDVVSTVGAGDSMVGGLIYGLLMRESSEHTLRLATAVA	282
WP_025798139.1	LGAEGALWVNASGAWLAKP-PACEVVSTVGAGDSMVGGLIYGLMMRESSDHTLRLATAVA	282
RDT52444.1	LGAEGALWVNASGEWIAKP-PSMEVVSTVGAGDSMVGGLIYGLLMRESSEHTLRLATAVA	282
WP_001653212.1	LGAEGALWVNASGEWIAKP-PSVDVVSTVGAGDSMVGGLIYGLLMRESSEHTLRLATAVA	282
EEZ4480454.1	LGAEGALWVNASGEWIAKP-PAVDVVSTVGAGDSMVGGLIYGLLMRESSEHTLRLATAVA	282
WP_163341112.1	LGAEGALWVNASGEWIAKP-PSVDVVSTVGAGDSMVGGLIYGLLMRESSEHTLRLATAVA	282

Ribokinase(P0A9J6)	AIAVTRKGAQPSVPWREEIDAFLDRQR-----	309
EHEC EDL933	ALAVSQSN-V-GITDRPQLAAMMARVDLQPFN	312
Uniprot_P0AEW9	ALAVSQSN-V-GITDRPQLAAMMARVDLQPFN	312
WP_025798139.1	ALAVSQSN-V-GISDRTQLAAMMARVDLKPFN	312
RDT52444.1	ALAVSQSN-V-GITDRTQLAAMMARVDLKPFN	312
WP_001653212.1	ALAVSQSN-V-GITDRPQLAAMMARVDLQPFN	312
EEZ4480454.1	ALAVSQSN-V-GITDRPQLAAMMARVDLQPFN	312
WP_163341112.1	ALAVSQSN-V-GITDRPQLAAMMARVDLQPFN	312
```

**Supplemental Figure 9. Increased ionic strength inhibits the Kinase Glo luciferase assay.**

No FruK was present in these control assays. (Top) Control assay with ATP only (no enzyme) shows that the luciferase assay could detect changes in this FruK product. (Bottom) Control assays with F-1,6-BP and ATP show that, if the increased ionic strength that results from increasing F-1,6-BP (x axis) or ATP (left versus right) concentrations is not balanced across the assay for the additional counter ions that accompany each metabolite, the luciferase activity is inhibited by the increasing ionic strength.

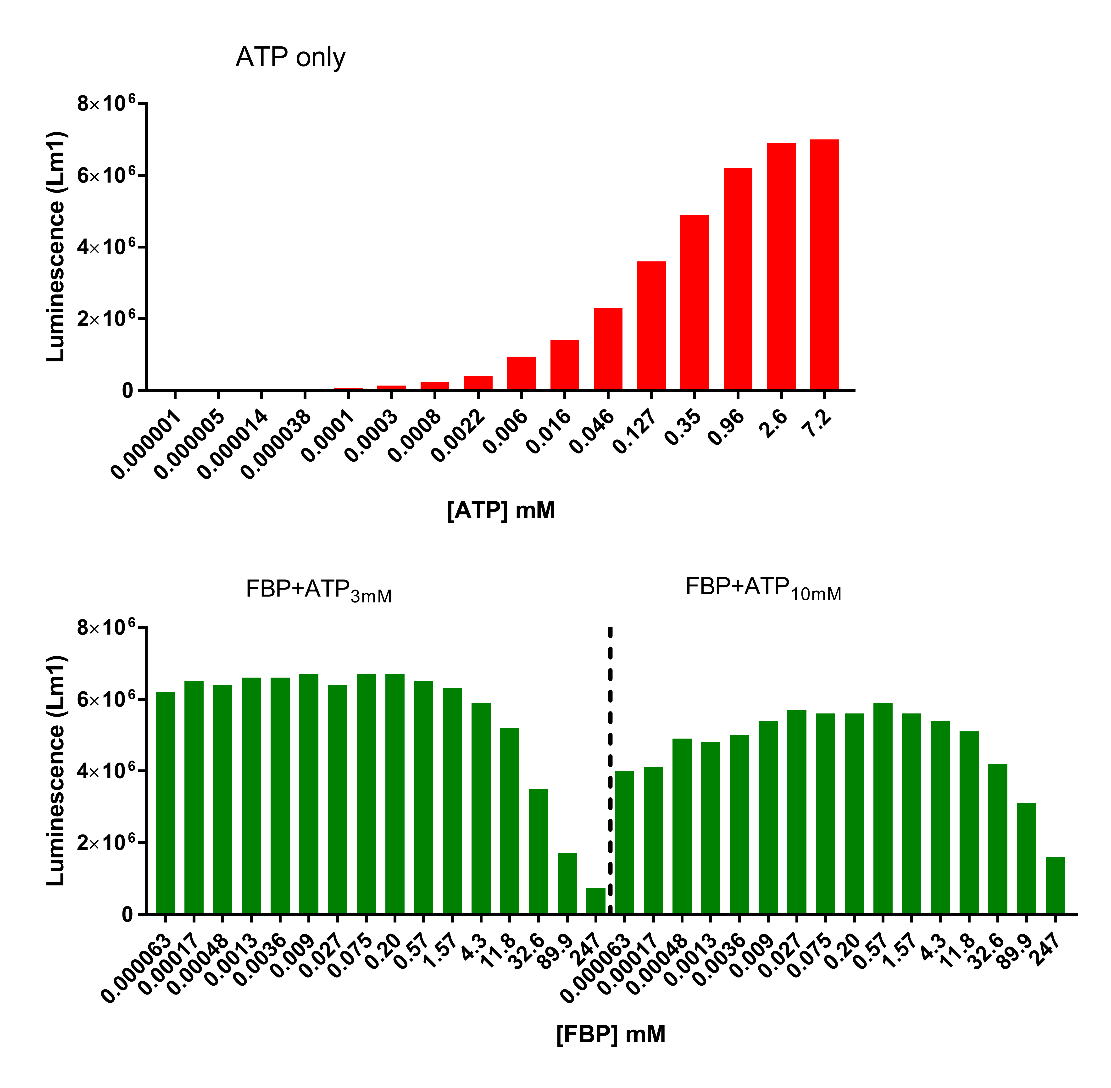

**Supplemental Methods: SDS-PAGE.** Once the PhastGel gradient gels used for initial studies were discontinued, we have not found any other commercial gels (even gradient gels spanning the same range) that resolve these two proteins. Fortunately, 15% gels following the original recipe of Laemmli^15^ can resolve Cra from FruK (see Figure 3 in the main text).

| ***SEPARATIVE GEL*** | | | | | | | | | |
| --- | --- | --- | --- | --- | --- | --- | --- | --- | --- |
|  | **Gel percentage** | | | | | | | | |
|  | **15%** | | **12.5%** | | | **10%** | | **6%** | |
| ***Component*** | ***4 gels*** | ***2 gels*** | ***4 gels*** | ***2 gels*** | | ***4 gels*** | ***2 gels*** | ***4 gels*** | ***2 gels*** |
| dH_2_O | 4.5 ml | 2.3 ml | 3.7 ml | 1.9 ml | | 5.4 ml | 2.7 ml | 8.0 ml | 4.0 ml |
| 30% Acrylamide mix | 10 ml | 5.0 ml | 8.4 ml | 4.2 ml | | 6.7 ml | 3.4 ml | 4.0 ml | 2.0 ml |
| Tris-HCl 1.0 M pH 8.8 | 5.1 ml | 2.6 ml | 7.5 ml | 3.8 ml | | 7.5 ml | 3.8 ml | 7.5 ml | 3.8 ml |
| 10% SDS (w/v) | 200 µl | 100 µl | 200 µl | 100 µl | | 200 µl | 100 µl | 200 µl | 100 µl |
| 10% APS | 200 µl | 100 µl | 200 µl | 100 µl | | 200 µl | 100 µl | 200 µl | 100 µl |
| TEMED | 8 µl | 4 µl | 12 µl | 6 µl | | 8 µl | 4 µl | 12 µl | 6 µl |
| **Total volume** | 20 ml | 10 ml | 20 ml | 10 ml | | 20 ml | 10 ml | 20 ml | 10 ml |
| **STACKING GEL** | | | | | | | | | |
|  | **Volume** | | | | | | | | |
|  | ***4 gels*** | | | | | ***2 gels*** | | | |
| dH_2_O | 6.85 ml | | | |  | 3.42 ml | | | |
| 30% Acrylamide mix | 1.7 ml | | | |  | 850 µl | | | |
| Tris-HCl 1.0 M pH 6.8 | 1.25 ml | | | |  | 625 µl | | | |
| 10% SDS (w/v) | 100 µl | | | |  | 50 µl | | | |
| 10% APS | 100 µl | | | |  | 50 µl | | | |
| TEMED | 10 µl | | | |  | 5 µl | | | |
| **Total volume** | 10 ml | | | |  | 5 ml | | | |

- 30% Acrylamide mix = 30 : 0.8 (Acrylamide : Methylenbisacrylamide)
- **10 X SDS-PAGE Running buffer** (pH 8.3):

Tris base	30.3 g

Glycine	144 g

10% SDS	100 ml

Add dH_2_O up to 1 L and store at room-temperature

- For 1 gel: 240 V; 20 mAmp. For 2 gels: 240 V; 40 mAmp

**Preparation of the protein sample:** Add 2X SDS Loading Buffer to the sample at a final 1X concentration; boil 3–5 min at 95°C; spin down briefly; load 10 – 20 µl on the SDS PAGE gel.

| ***2 X SDS Loading Buffer ( 10 ml)*** | | | |
| --- | --- | --- | --- |
|  |  |  | ***Final concentration*** |
| 1.0 M Tris-HCl pH 6.8 | 1.2 ml |  | 60 mM |
| 10% SDS | 2.0 ml |  | 1% |
| β-mercaptoethanol | 200 µl |  | 1% |
| 50% Glycerol | 4.0 ml |  | 20% |
| Bromphenol Blue | 0.001 g |  | 0.01% |
| **dH_2_O** | **Up to 10 ml** |  |  |

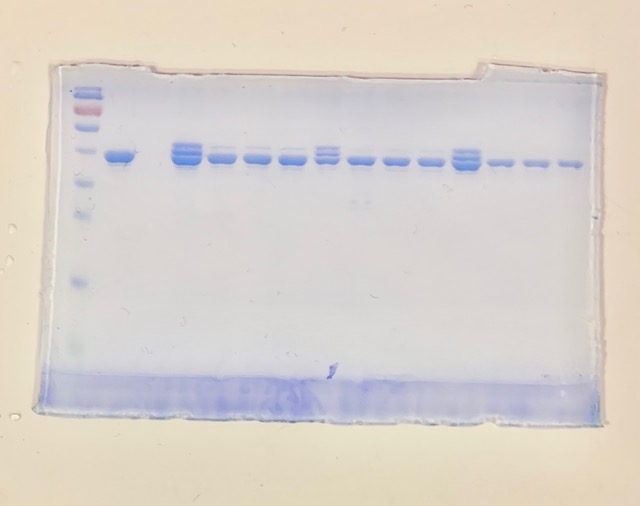

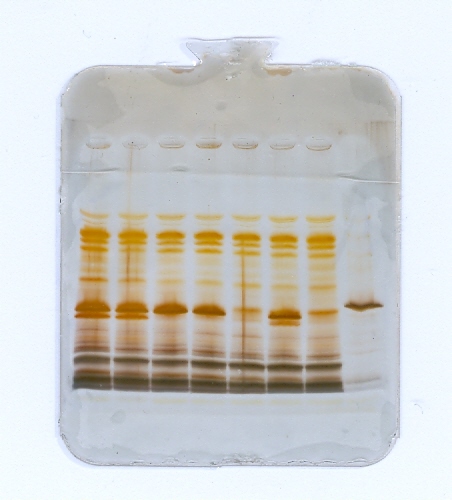
**Supplemental Figure 10. Uncropped gel images for Figure 3 in the main text.**

**B**

**A**

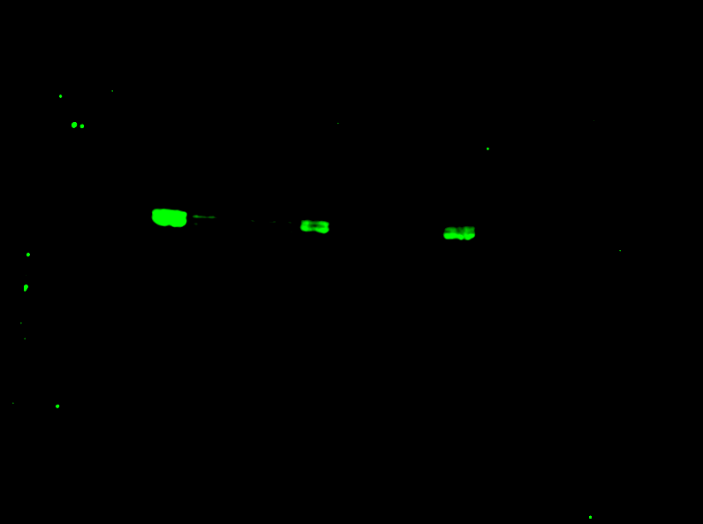

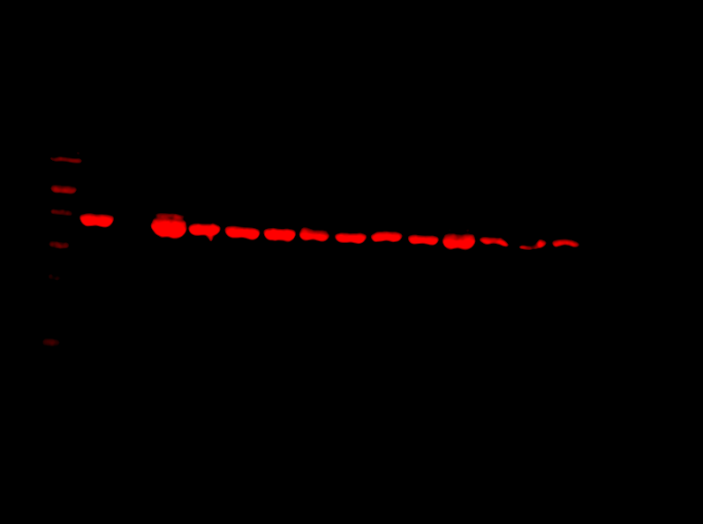

**C**

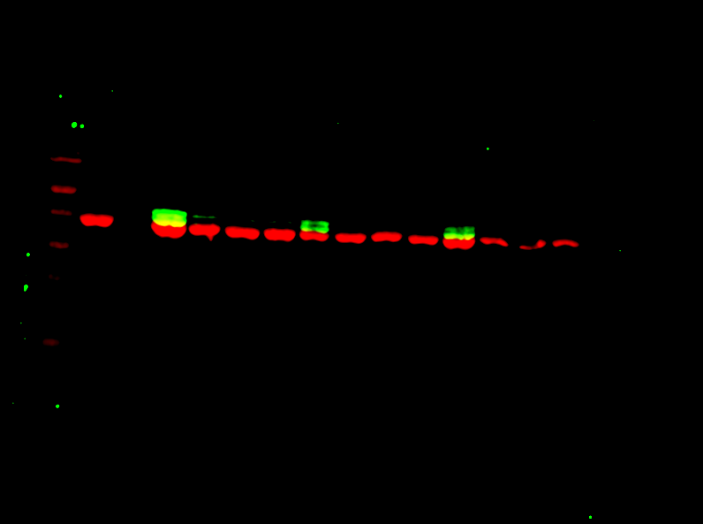

In the gels shown in B/C, the first four lanes are in Figure 3 of the main text. The remaining lanes show replicates or contained samples that are not relevant to the results reported in this manuscript.
